# Intercellular communication in the brain via dendritic nanotubular network

**DOI:** 10.1101/2025.05.20.655147

**Authors:** Minhyeok Chang, Sarah Krüssel, Laxmi Kumar Parajuli, Juhyun Kim, Daniel Lee, Alec Merodio, Jaeyoung Kwon, Shigeo Okabe, Hyung-Bae Kwon

## Abstract

Recent studies have identified intercellular networks for material exchange by bridge-like nanotubular structures, yet their existence in neurons remains unexplored within the brain. Here, we identified long, thin dendritic filopodia that establish direct dendrite-to-dendrite contacts, forming dendritic nanotubes (DNTs) in mammalian brains. Using super-resolution microscopy, we characterized their unique molecular composition and dynamics in dissociated neurons, enabling Ca^2+^ propagation over distances. Utilizing imaging and machine-learning-based analysis, we confirmed the *in situ* presence of DNTs connecting dendrites to other dendrites whose anatomical features are distinguished from synaptic dendritic spines. DNTs mediate the active transport of small molecules or human amyloid-beta (Aβ), implicating the role of DNT network in AD pathology. Notably, DNT levels increased prior to the onset of amyloid plaque deposits in the mPFC of APP/PS1 mice. Computational simulations predicted the progression of amyloidosis, providing insight into the mechanisms underlying neurodegeneration through these DNTs. This study unveils a previously unrecognized nanotubular network, highlighting another dimension of neuronal connectivity beyond synapses.

## Introduction

Over the last 20 years, advances in microscopy have unveiled noncanonical cellular structures that bridge distant cells. These structures are now recognized as a common aspect of cellular biology, facilitating specialized remote communication, by transporting ‘cargoes’ to exclusively connected cells. This long-range, specific signaling contrasts with traditional pathways such as diffusion-based signal release or gap junction-mediated exchange which is limited to adjacent cells. For instance, developing cells produce cytonemes to transfer morphogens to specific target regions (*1, 2*). *In vitro* studies of immortalized cells have identified tunneling nanotubes (TNTs) connecting cellular bodies via membrane-fused terminals (*3*). TNTs serve as general routes for intercellular material exchange – ranging from ions to organelles (*4*) - with recent studies confirm their presence *in vivo* (*5, 6*). Despite variations in terminology, these connections share a common morphology: an intricate nanotubular architecture, only hundreds of nanometers thick but extending for micrometers. It has been proposed that these nanotubular connections originate from filopodia, thin membranous protrusions (*7, 8*), as recently supported by their formation process (*9*).

Do neurons form nanotubular connections in the brain? Previous studies have claimed the presence of soma-to-soma TNTs in dissociated neurons of early DIVs (*10-13*), similar to those observed in immortalized non-neuronal cells. However, such connections are unlikely to be prevalent in the mature brain, given the known mechanisms of nanotubular formation by filopodia (*9*) or contact between cell bodies (*3*). Neuronal somas are neither mobile nor present filopodia after early developmental stages (*14*). In contrast, dendrites develop various protrusions, including dendritic filopodia, primarily studied as precursors to synaptogenesis (*7, 15*) by contacting presynaptic partners. Evidence suggests unconventional roles for dendritic filopodia in different contexts, supported by stable filopodia independent of spine formation (*16, 17*). Our hypothesis posits that dendritic filopodia can conditionally develop into bridge-like structures distinct from previously described synaptic connections or dendritic branches.

Recent studies have suggested the spreading of proteins via nanotubular propagation between astrocytes and neurons (*18*) or between neurons of different brain regions (*19*). However, the nature of this transport remains unclear, as the presumed nanotubes have not been distinctly characterized morphologically or molecularly and have only been assumed to be nanotubes. Given the strong link between cell-to-cell spreading and aggregation of pathological peptides, this potential nanotubular connectivity in the brain may contribute to neurodegeneration including Alzheimer’s disease (AD) (*20-22*). Although senile plaques are a significant hallmark of AD progression, the precise mechanism of beta-amyloid (Aβ) in AD pathology has remained controversial for decades. Consequently, efforts to diagnose and treat the disease by targeting extracellular Aβ have struggled (*23*). In this context, some researchers have focused on intracellular Aβ, which appears in early AD (*24-26*) before plaque deposition. Remarkably, microinjected oligomeric Aβ has been reported to transfer between primary hippocampal rat neurons via direct cellular connections through an unknown nonsynaptic mechanism (*27*), suggesting nanotubular connectivity.

Altogether, the nanoscale network in the brain seems to play a pivotal role in normal brain functions or brain pathology. However, the minute physical sizes, dynamic nature, absence of markers, and their embedding within immense neuronal connectivity have made it challenging to investigate nanotubular connections among neurons. To reveal this unseen network, we used a specialized microscopy technique called dSRRF (super-resolution radial fluctuation microscopy aided by volumetric deconvolution) combined with an optimized sample preparation technique utilizing mouse brain tissue clearing. This technical advancement enables detailed examination of dendritic protrusions at the nanoscale. Our results indicate that a nanotubular network originating from dendritic filopodia facilitates the propagation of calcium and molecules, including Aβ, suggesting a novel aspect of amyloid pathology in AD. Cortical pyramidal neurons are interconnected through this structure, and this connectivity is elevated in the medial prefrontal cortex (mPFC) of APP/PS1 mice brains at the early pathological stage of 3 months. Further investigation revealed how nanotubular connections contribute to selective intraneuronal accumulation of Aβ. This study provides evidence of long-range neuronal connectivity beyond canonical synapses.

## Results

### Nanoscale dendritic filopodial bridges in mice and human

Nanoscale filopodia embedding within a tissue are difficult to resolve with a diffraction-limited microscope, and even when stochastic fluorescent tagging facilitates their detection, the sparseness makes it difficult to determine what cellular compartments they are contacting. Recent advances in volumetric electron microscopy (EM) imaging have allowed the clear visualization of dendritic filopodial morphology at nanometer-scaled voxels. We first reanalyze the previously published EM data to investigate the existence of filopodial bridges in neurons and visualize where they originate from and end. This analysis was completely separate and unbiased because the original experimental designs of these studies were not intended to study nanotubular dendritic filopodia. We examined two pre-acquired EM datasets of dendritic morphology: the somatosensory cortical layer 1 of the mouse brain (**Supplementary Fig. 1**)(*28*) and the cortical layers 2 and 5 of the middle temporal gyrus in the human brain (H01 dataset; **Supplementary Figs. 2, 3**)(*29*). We discovered that certain dendritic filopodia in both datasets formed membrane-membrane contacts with neighboring dendrites but without the postsynaptic density typically found in excitatory synapses (**Supplementary Figs. 1, 2A-D, 3, and Supplementary Movies 1-3**). The absence of chemical synapse features on these filopodia was confirmed by the annotations provided for synapses in the H01 dataset (**Supplementary Fig. 2A-D**; Methods). A notable portion of nonsynaptic dendritic filopodia (thickness of heads < 400 nm) established closed ended membrane-membrane contacts with other dendrites, significantly from long (end-to-end distance > 3 μm) filopodia, while synaptic spines represented less chance of dendritic contact despite of their larger head sizes (**Supplementary Fig. 2E-M**). The dendritic contacts particularly located at or near the filopodial tips (**Supplementary Fig. 2N**). We did not observe any open ended tunneling by membrane-membrane fusion in dendrites. Additionally, we did not find any structures connecting soma-to-soma directly. These structures are distinguished from previously reported TNTs that normally connect cell bodies.

Given that these newly discovered structures neither connect cell bodies nor possess open or tunneled terminals like TNTs, we refer to them as dendritic nanotubes (DNTs) hereafter.

### Molecular characteristics of dendritic nanotubes in dissociated cortical neurons

Our investigation in EM images revealed the potential presence of DNTs connecting dendrites in the brain. However, further systematic characterization of their structure and molecular compositions was needed for a better understanding of their physiology. We modified SRRF (super-resolution radial fluctuation microscopy) (*30*), a versatile super resolution approach, for volumetric imaging with a preceding deconvolution process (dSRRF; **Supplementary Fig. 4**, Methods). Using dSRRF, we resolved neuronal structures at the nanoscale to characterize filopodia-forming DNTs in the dissociated culture of cortical neurons. Super-resolution imaging of live mouse cortical neurons (DIV9), stained with neuronal-membrane specific dye (NeuO), revealed membranous bridge-like structures between neurites (**Fig. 1A**; Methods). These intermediate extensions (interextensions) may be segments of thin axonal neurites that are developing branches or traversing hollow spaces. However, some of these extensions did not contain tubulin, which would typically be present in such structures (**Fig. 1B**). Certain fine neurites may show reduced fluorescence signals from tubulin due to diminished inner contents. We compared the fluorescent signals of tubulin-negative and tubulin-positive structures after normalization by width (**Fig. 1C**). Tubulin-negative interextensions still exhibited distinct fluorescent signal strengths after normalization (**Fig. 1C**) and width (**Fig. 1D**), suggesting these structures are likely dendritic nanotubes.

**Figure 1.**
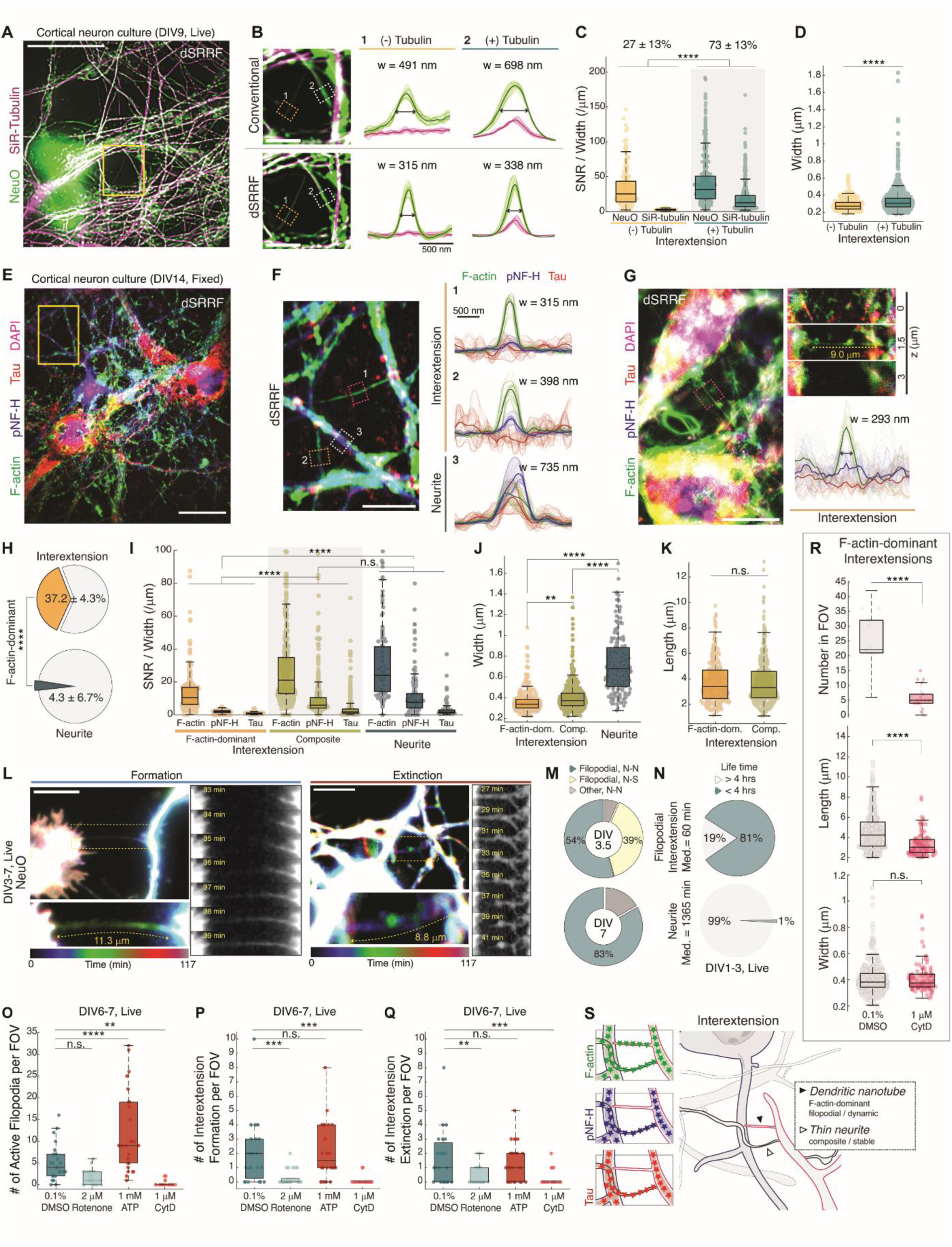
Dendritic nanotubes (DNTs) in dissociated cortical neurons. **(A)** Representative image of live cortical neurons in dissociated culture (DIV9) by dSRRF microscopy which visualizes neuronal membrane (NeuO, green) and microtubule (SiR-tubulin, magenta). n_volume_ = 100 for dSRRF reconstruction. **(B)** Interextensions between two neurites (branched directly from somas or other neurites) shown in conventional (top) and dSRRF (bottom) microscopies, magnified from the boxed region (yellow) in (A). Cross-sectional intensity profiles averaged from colored boxes (left) show distinct cytoskeletal composition between interextensions. **(C)** Signal-to-noise ratio (SNR) of averaged intensity profile along structures, which normalized by width (FWHM; Full Width at Half Maximum) for each cytoskeletal component from tubulin-negative (left, n = 230; SNR_tubulin_ < 1.2) and tubulin-positive (right, n = 995; SNR_tubulin_ ≥ 1.2) interextensions. *P* = 2.9 × 10^-26^ between groups by unbalanced two-way ANOVA. The fraction of each group was represented (top, mean ± s.d.; n_FOV_ = 59). **(D)** Width of structures (n = 230 and 995 for (-) and (+) tubulin; *P* = 1.6 × 10^-13^ by two-tailed unequal variances t-test). **(E)** Representative multicolor dSRRF image of fixed cortical neurons in dissociated culture (DIV14) with staining of three cytoskeletal proteins of F-actin (green), neurofilament (pNF-H, blue), and tau (red). n_volume_ = 1,000 for dSRRF reconstruction. **(F)** Interextensions between two neurites in dSRRF microscopies, magnified from the boxed region (yellow) in (E), and their cross-sectional intensity profiles. **(G)** Another example of an interextension linking a soma and a neurite (left), its axial montage (right, top), and cross-sectional intensity profile (right, bottom). **(H)** Percentage of F-actin-only structures in interextensions and neurites (n_FOV_ = 11) whose SNR for pNF-H and tau < 1.2 (two-tailed unequal variances t-test, *P* = 1.2 × 10^-10^). **(I)** SNR of averaged intensity profile along structures, normalized by width for each cytoskeletal component from F-actin-only intermextensions (left, n = 302), composite interextensions (middle, n = 452), and neurites (right, n = 191). *P* = 8.9 × 10^-12^ for Actin-IB vs. Comp-IB, *P* = 1.7 × 10^-9^ for Actin-IB vs. Primary, and *P* = 0.80 for Comp-IB vs. Primary by unbalanced two-way ANOVA, *P* = 2.0 × 10^-13^ among groups. **(J-K)** Width and length of structures (*P* = 0.0036 for Actin-IB vs. Comp-IB, *P* = 0 for Actin-IB vs. Primary, and *P* = 0 for Comp-IB vs. Primary by one-way ANOVA; *P* = 1.9 × 10^-98^ for (J), *P* = 0.95 for Actin-IB vs. Comp-IB by two-tailed unequal variances t-test for (K)). **(L)** Representative images displaying formation (top) and extinction (bottom) of interextensions from and to filopodia observed in time-lapse imaging of NeuO-stained neurons for 2 hours (video rate = 1 min). **(M)** Percentages of interextension formation connecting neurites (N-N) or a neurite and a soma (N-S) by filopodial contact or another mechanism (i.e. shaft contact) on DIV3.5 and DIV7 (n_exp_ = 37 and 35; n_event_ = 46 and 18; Filopodial formation = 93 % and 83 % for DIV3.5 and DIV7 respectively). **(N)** Temporal stability of interextensions (top; n = 59) and primary neurites (bottom; n = 73) is shown by percentages of structures presented longer or shorter than 4 hours in time-lapse imaging (66 hours, video rate = 15 min) of cortical neurons stained by 100 nM SiR-actin. **(O-Q)** The pharmacological effects of interextension formation from filopodia in time lapse imaging (1-hour-long, video rate = 1 min). The numbers of active filopodia (O), formation events (P), and extinction events (Q) were counted in the FOV (87 × 87 μm^2^) after 60-min-long incubation with 0.1% DMSO (n_FOV_ = 23), 2 μM Rotenone (n_FOV_ = 25; for intracellular ATP depletion by inhibiting mitochondrial complex I), 1 mM ATP (n_FOV_ = 26; for ATP supplement), and 1 μM Cytochalasin-D (n_FOV_ = 26; for inhibition of actin filament polymerization). *P* = 0.13 for DMSO vs. Rotenone, *P* = 7.1 × 10^-5^ for DMSO vs. ATP, and *P* = 0.009 for DMSO vs. Cyt-D by one-way ANOVA; *P* = 1.7 × 10^-12^ for (O). *P* = 0.0007 for DMSO vs. Rotenone, *P* = 1 for DMSO vs. ATP, and *P* = 0.0001 for DMSO vs. Cyt-D by one-way ANOVA; *P* = 2.5 × 10^-7^ for (P). *P* = 0.002 for DMSO vs. Rotenone, *P* = 1 for DMSO vs. ATP, and *P* = 0.0003 for DMSO vs. Cyt-D by one-way ANOVA; *P* = 2.7 × 10^-5^ for (Q). **(R)** The effect of 1 μM Cytochalasin-D (Cyt-D) on the number in the FOV (87 × 87 μm^2^), length, and width of F-actin-dominant interextensions or dendritic nanotubes (incubation for 90 min). n_FOV_ = 33 and 18, n = 722 and 107 for DMSO (control) and Cyt-D. *P* = 2.5 × 10^-10^ for DMSO vs. Cyt-D by One-way ANOVA (*P* = 1.6 × 10^-12^) for normalized density; *P* = 1.3 × 10^-10^ for DMSO vs. Cyt-D, *P* = 0.0001 by One-way ANOVA (*P* = 4.7 × 10^-15^) for length; *P* = 1 for DMSO vs. Cyt-D by One-way ANOVA (*P* = 0.38) for width. **(S)** Illustration of a dendritic nanotube or F-actin dominant interextension as a distinct structure from a composite interextension, possibly a part of thin neurites. n.s., not significant; * *P* < 0.05; ** *P* < 0.01; *** *P* < 0.001; **** *P* < 0.0001; In the box plots, the midline, box size, and whisker indicate median, 25-75^th^ percentile, and ±2.7σ, respectively. Bonferroni correction followed all the ANOVA tests. Scale bars = 20 μm for (A), 10 μm for (E), (G), (L), and 5 μm for (B), (F). Widths of (B), (F), and (G) were defined by the full-width-half-maximum (FWHM) of F-actin signal profiles.

To investigate the identity of these interextensions as nanotubes, we used multi-color dSRRF imaging of fixed dissociated cortical neurons (DIV14) to detect filamentous actin (F-actin), neurofilament (pNF-H), and tau on these structures (**Fig. 1E**; Methods). F-actin emerged as a primary component of filopodia and nanotubular connections in previous literature (*3, 4, 9*). To preserve fragile nanostructures, we employed a cytoskeleton-preserving fixation protocol (*31*). We consistently observed interextensions with dominant F-actin but no pNF-H or Tau signals, connecting neurites (**Fig. 1F**) or rarely neurites and somas (**Fig. 1G**; 37% in total, **Fig. 1H**). Normalized fluorescence signals from F-actin-dominant interextensions revealed their distinct molecular compositions, differing from the shared contents of neurites and composite interextensions – probably thin neurites (**Fig. 1I**). These two groups of interextensions were indistinguishable without molecular characterization due to negligible differences in their widths (335 nm vs 368 nm, **Fig. 1J**) or lengths (3.4 μm vs 3.3 μm, **Fig. 1K**).

To confirm whether these are indeed nanotubular structures that originate from filopodia, we attempted to observe their real-time formation and elimination processes. For convenience of imaging, we used neurons with less projections before DIV7 (**Supplementary Fig. 6A-D**). Because we did not predict where the filopodium would grow, we developed an automated long-term imaging system that enables to take imaging of multiple region-of-interests over a long period of time at short time intervals. 2-hour-long time-lapse imaging (video rate = 1 min) successfully captured several cases of the formation and elimination of extensions in real-time (**Fig. 1L, Supplementary Movies 4 and 5**; Methods; the average speeds of formation and extinction = 0.62 and 0.77 μm/min, **Supplementary Fig. 6E**). The formation of neurite-soma interextensions was frequent as neurite-neurite interextensions on DIV3.5 but significantly decreased by DIV7 (**Fig. 1M and Supplementary Fig. 6F**). To measure temporal stability of the extensions, we attempted imaging for daily scale (66 hrs) with increased time interval to minimize photobleaching (video rate = 15 min). The life span of extensions was shorter compared to stable thicker neurites (**Fig. 1N**).

Dynamic nature of these filopodial extensions implies metabolic regulation. Intracellular ATP depletion by Rotenone inhibited formation and elimination of the interextensions, but elevated activity of filopodia by ATP supplement did not promote the processes (**Fig. 1O-Q**).

Therefore, ATP-regulating dynamics is necessary but not sufficient for filopodial interextension formation. Cytochalasin-D (Cyt-D), an inhibitor of actin polymerization shown to eliminate dendritic filopodia (*32*) but preserve elongation of neurites (*33*) annihilated filopodial activity and reduced both the appearance and length of F-actin-dominant interextensions, suggesting it as an efficient inhibitor (**Fig. 1O-R**; Methods). Overall, we conclude that F-actin-dominant interextensions are potential DNTs connecting neurons and their structure is dynamically regulated in dissociated cultures of cortical neurons (**Fig. 1S**).

### Calcium transfer via dendritic nanotubes in cortical neurons

We and others have shown calcium transfer through TNTs connecting cellular bodies of HeLa (*9, 34*) or pericytes (*5*). As previously reported (*10-13*), we employed a similar experimental approach to investigate calcium propagation through the somatic nanotubes. By uncaging caged calcium (DMNPE-4-AM) in a soma using focused ultraviolet light (λ = 365 nm), we observed increased calcium indicator signals (Cal-590-AM) along a somatic nanotube and in the connected neighboring soma (**Fig. 2A-C, Supplementary Movie 6 and 7**). This transfer through somatic nanotubes prompted further investigation into the capability of dendritic nanotubes to facilitate calcium propagation over longer distances within neuronal networks. To explore this, we monitored intensity changes in neighboring neurons following UV-targeting of a single neuron from cultures ranging from DIV13-17 for 50-sec-long UV exposure, which saturated indicators in the target (**Fig. 2D**; Methods). Neurons within the field of view (FOV) exhibited heterogeneous changes in indicator intensities, supporting cell-specific calcium propagation rather than experimental artifacts (**Fig. 2E-G**).

**Figure 2.**
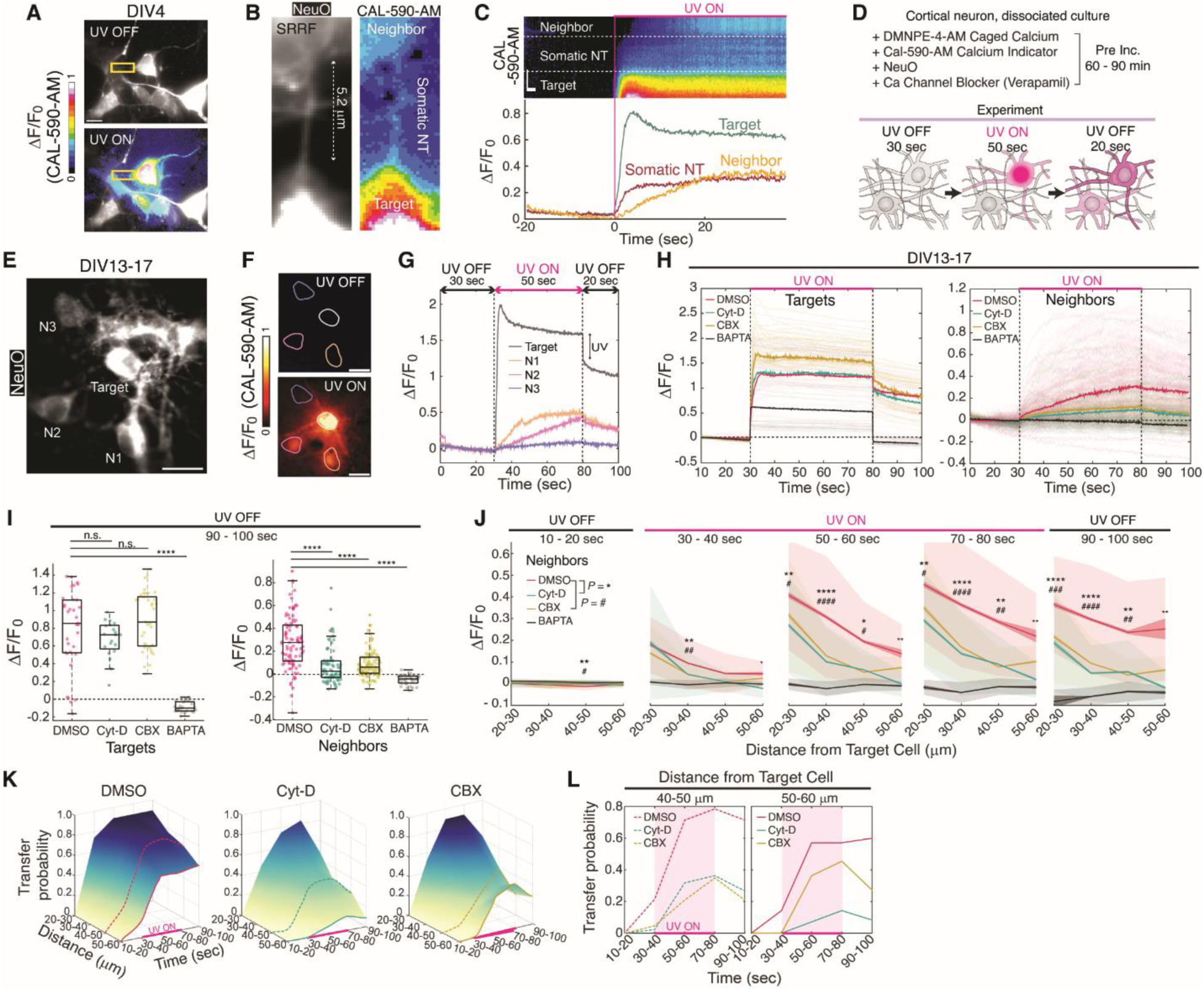
Ca^2+^ transfer by DNT between distant neurons in dissociated culture. **(A)** An example showing the transfer of uncaged calcium between cortical neurons (DIV4). Exclusive UV (λ = 365 nm) exposure on a single neuron induced immediate uncaging of DMNPE-4-AM and the rapid rise of Ca^2+^ concentration in the targeted cell where NeuO stained neurons. **(B-C)** Somatic nanotube-mediated calcium propagation between neurons and its time trace. **(D)** Schematic of the experimental procedure to detect calcium transfer between cortical neurons in culture (DIV13-17). For each experimental session, the Cal-590-AM indicator intensities in the UV-exposed neuron (*target*) and surrounding neurons (*neighbors*) were monitored for 100 seconds. More details in Methods. **(E)** Representative image from an experimental session showing the locations of a *target* and three *neighbors*. **(F)** Images of the calcium indicator signals (ΔF/F_0_) before and 30 seconds after UV exposure on the *target* neuron. **(G)** The time traces of the calcium indicator intensity from the *target* and *neighboring* neurons display distinct signal responses. A sudden drop when UV was off in the target trace was not observed in the *neighbors*, indicating no direct UV exposure was delivered to them. **(H)** Time traces of Cal-590-AM intensity from *targets* (n = 34; top) and *neighbors* (n = 93; bottom) in the control group (+0.1% DMSO for pre-incubation); *targets* (n = 34; top) and *neighbors* (n = 122; bottom) in the DNT reduced group (+1 μM Cytochalasin-D for pre-incubation); *targets* (n = 37; top) and *neighbors* (n = 129; bottom) in the connexin-blocked group (+100 μM Carbenoxolone for pre-incubation); and *targets* (n = 30; top) and *neighbors* (n = 43; bottom) in the intracellular calcium-chelated group (+50 μM BAPTA-AM for pre-incubation). Thick lines represent the median of individual traces. **(I)** Averaged calcium-indicator intensity in *target* neurons (n = 34 for the control group; n = 34 for the Cyt-D group; n = 37 for the CBX group; n = 30 for the BAPTA group) and *neighbor* neurons (n = 93 for the control group; n = 122 for the Cyt-D group; n = 129 for the CBX group; n = 43 for the BAPTA group) after UV exposure in (H) (t = 90 – 100 sec; One-way ANOVA with Bonferroni correction for comparison, *P* = 1.2 × 10^-21^ and 1.7 × 10^-24^ for *targets* and *neighbors*; details in Supplementary Table). **(J)** Calcium signals in *neighbors* represented by the distance from *targets.* The intracellular signals were averaged for the given time intervals and grouped by the distance (bin width = 5 μm; line = the mean of each bin; bright shade = the standard error in each bin; dark shade = the standard deviation in each bin; One-way ANOVA with Bonferroni correction to compare experimental groups; P-values are displayed by the numbers of asterisks for DMSO vs. Cyt-D and hashes for DMSO vs. CBX; details in Supplementary Table). **(K)** Transfer probability (the proportion of *neighbors* showing SNR change > 0.1) in the control group, the Cyt-D treated group, and the CBX-treated group regarding time and distance from target cells. Surface plots were generated from the bins in (J). Details in Methods. **(L)** Comparison of transfer probabilities in long distances by time (left, 40 – 50 μm; right, 50 -60 μm). n.s., not significant; * *P* < 0.05; ** *P* < 0.01; *** *P* < 0.001; **** *P* < 0.0001; In the box plots, the midline, box size, and whisker indicate median, 25-75^th^ percentile, and ±2.7σ, respectively. Scale bars = 10 μm.

To verify that the observed propagation was DNT-mediated, we analyzed indicator intensities in neighboring neurons under four pre-incubation conditions; DMSO, Cyt-D, Carbenoxolone (CBX) (*35*), and BAPTA-AM (**Fig. 2H**). Reduction of DNTs by Cyt-D significantly attenuated the transferred calcium signal in neighbors compared to the control group (DMSO), strongly suggesting DNT involvement in the transfer mechanism (**Fig. 2I**). Previous studies (*5, 36*) have proposed the involvement of gap junctions in TNT-mediated calcium propagation. Interestingly, our data suggests that connexin is also involved in DNT-mediated calcium propagation; neurons treated with CBX, a connexin blocker, exhibited decreased calcium transfer to neighboring cells (**Fig. 2I**). On the contrary, pharmacological inhibition of other calcium sources such as N-methyl-D-aspartate (NMDA) receptor, L-type calcium channel, and pannexin-1 did not suppress the transfer (**Supplementary Fig. 7**).

This condition-dependent reduction in transfer was observed across all distances up to 60 μm, the maximum distance within our FOV (**Fig. 2J**). For a comprehensive analysis, we compared the transfer probabilities in the neighborhoods of experimental groups based on distance from the target neurons and experimental time (**Fig. 2K**). Notably, DNT reduction and connexin blocking did not yield identical results; while DNT reduction led to a more globally diminished propagation, particularly over long distances (**Fig. 2L**), suggesting a gap-junction-independent mechanism of connection through DNTs.

### Morphological characterization of *in situ* dendritic nanotubes in mouse brain

The superior resolution of EM successfully revealed the existence of DNTs in mice and human brains. However, to gain a deeper understanding of the physiology of DNT *in situ*, a more accessible approach was required. We attempted optical imaging of Thy1-GFP-M (*37*) mouse brains, which exhibits preferential GFP expression in sparsely scattered neurons within deep cortical layers. Imaging the layer 5 visual cortex of a Thy1-GFP-M mouse brain after tissue clearing (CUBIC-RIMS) (*38, 39*) with light sheet microscopy, we resolved DNT-like structures between dendrites of different neurons (**Fig. 3A-C**). However, limited resolution restricted the precise characterization of their morphology, which was expected to be essentially similar to DNTs in dissociated cultures (**Fig. 3D**, **Fig. 1F**).

**Figure 3.**
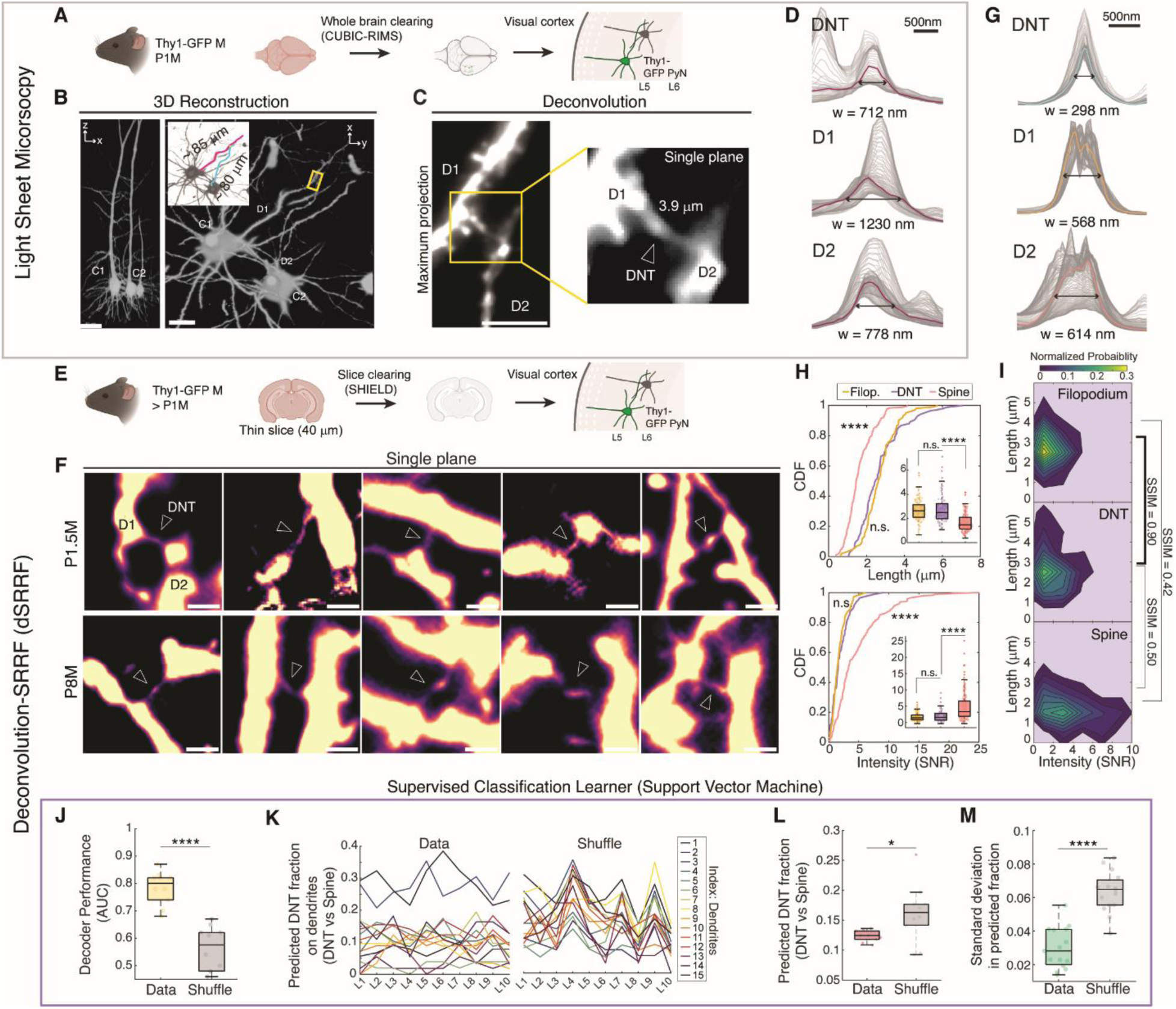
*in situ* DNTs of cortical neurons in mouse brain. **(A)** Schematic of sample preparation to image DNTs in tissue-cleared whole mouse brain (Thy1-GFP-M) using light sheet microscopy. **(B-C)** Representative image of a DNT connecting two basal dendrites (D1, D2) from layer 5 pyramidal neurons (C1, C2) in the visual cortex. Arrow indicates the position of DNT. **(D)** Cross-sectional intensity profiles for the DNT and two connected dendrites. Widths are FWHMs of the median intensity profiles. **(E)** Schematic of sample preparation for DNT imaging in tissue-cleared mouse brain slices by dSRRF. **(F)** Representative dSRRF images of DNTs observed in the layer 5 visual cortices of P1.5M brains (top) and P8M brains (bottom). Arrow = position of DNT. **(G)** Cross-sectional intensity profiles for the DNT and two connected dendrites (D1, D2) from (F). FWHMs from the median intensity profiles were measured for the widths of the structures. **(H)** CDFs for lengths and median intensities of dendritic filopodia (Filop., n = 103), dendrite-dendrite DNTs (n = 83), and axon-contacting dendritic spines (n = 176) from P8M brains (n_mouse_ = 2). Two-sampled Kolmogorov-Smirnov test was performed for DNT vs. Spine (*P* = 1.6 xyx 10^-12^ for top; *P* = 1.8 × 10^-7^ for bottom) and DNT vs. Filop. (*P* = 0.36 for top; *P* = 0.61 for bottom). (Inlets) Multi-comparison of the structural groups by One-way ANOVA with Bonferroni correction (*P* = 1.4 × 10^-24^ for top; *P* = 1.0 × 10^-16^ for bottom; (top) *P* = 2.0 × 10^-17^ for DNT vs. Spine and *P* = 1 for DNT vs. Filop.; (bottom) *P* = 1.4 × 10^-10^ for DNT vs. Spine and *P* = 1 for DNT vs. Filop.). **(I)** Heat map presentation of normalized probability for lengths and median intensities of structural groups. The structural similarity index (SSIM) was measured to quantify the similarity of histograms. **(J)** Decoder performance of supervised classification learner (quadratic SVM) for DNT / Spine classification. Ten models were trained with 22 morphological features of pre-labeled structures (n = 259) each for the actual data set (Data) and shuffled data set (Shuffle) and compared. *P* = 1.6 × 10^-6^ by two-tailed t-test with equal variances. **(K)** Data and Shuffle learners predicted the fraction of DNTs from 940 unlabeled dendritic protrusions. *P* = 0.026 by two-tailed equal variances t-test. **(L)** Predicted DNT fraction on dendrites by Data and Shuffle learners (n_dendrite_ = 15, n_learner_ = 10). **(M)** Standard deviation in the prediction of DNT fraction on each dendrite by learners. (n_dendrite_ = 15; *P* = 3.5 × 10^-8^ by two-tailed t-test with equal variances). n.s., not significant; * *P* < 0.05; **** *P* < 0.0001; In the box plots, the midline, box size, and whisker indicate median, 25-75^th^ percentile, and ±2.7σ, respectively. Scale bars = 50 μm (B, left), 20 μm (B, right), 5 μm (C), and 1 μm (F).

We optimized dSRRF imaging of dendritic structures from brain slices by minimizing undesired fluorescence from the background through tissue-clearing and thin slicing (**Fig. 3E**, **Supplementary Figs. 4 and 5**). Leveraging the fast-acquisition speed of wide-field microscopy, we acquired multiple volumetric stacks for SRRF reconstruction in a relatively short time (∼ 4 min for 65 × 65 × 40 μm^3^ by 500 nm axial step when N_volume_ = 100). Super-resolved images of layer 5 pyramidal neurons revealed DNTs as thin bridge structures between two dendrites, observed in both young (P1.5M) and adult (P8M) mice (**Fig. 3F**), exhibiting similar morphology to DNTs in culture (**Fig. 3G**, **Fig. 1F**) and EM-resolved DNTs (**Supplementary Fig. 2**). The distribution of two morphological features - length and median intensity in SNR along the structure - appeared similar between presumed DNTs and dendritic filopodia (thin protrusions whose terminal contacts were unresolved due to sparse GFP expression), whereas dendritic spines (protrusions contacting axons) displayed significantly distinct distribution (**Fig. 3H**). This distinction was more evident when comparing features in two-dimensional representation; the structural similarity index (SSIM) indicated high similarity of DNTs and filopodia, but not between DNTs and spines (**Fig. 3I**).

Although DNTs are statistically distinct from spines, it was challenging to discriminate between the two structures using simple numeric thresholds for just two features, likely due to the diverse shapes of dendritic spines (**Supplementary Fig. 8A**). We devised a classification method for dendritic protrusions training supported vector machine (SVM) learners under supervision (**Supplementary Fig. 8B**; Methods). For accurate training, we provided 22 unbiased morphological features with labels to the learners. Data-trained learners (Data-learner) exhibited significantly enhanced performance compared to learners trained with shuffled labels (Shuffle-learner; **Fig. 3J, Supplementary Fig. 8C**), implying a clear morphological distinction for DNTs. The path length was the most significant but not the sole distinctive feature (**Supplementary Fig. 8D-E**). Using these learners, we classified 940 unlabeled dendritic protrusions from 15 dendrites into either DNT or Spine groups (**Fig. 3K**). The fraction of DNTs was predicted to be 12% by 10 Data-learners which significantly differed from the prediction by 10 Shuffle-learners (**Fig. 3L**; Discussion). Data-learners predicted consistent fractions of DNTs for individual dendrites, whereas Shuffle-learners produced inconsistent results (**Fig. 3M**), supporting the relevance of characterization by trained learners.

### Dendritic nanotube mediates Aβ propagation between distant neurons in mPFC

In our study, newly discovered DNTs are proposed as the underlying mechanism for molecular propagation in the brain. To verify whether DNTs mediate small molecule transfer to neighboring neurons, we infused fluorescent dyes through whole-cell patch clamping in acutely prepared mouse brain slices (Methods). Two different molecular weights (0.7 kDa and 10 kDa) emitting different fluorescent colors were mixed and loaded into the same patch clamp pipette to determine if DNT-mediated transfer depends on molecular size. We detected both dye molecules were transferred to several cells near the patched target neuron (**Supplementary Fig. 9A-C**). The high correlation of the two dye molecules suggested a shared mechanism of transfer (**Supplementary Fig. 9D**). It also suggests that DNTs mediate the propagation of molecules up to 10 kDa. Notably, the transfer was heterogeneous and most prominent near the dendrites of the patched cell (**Supplementary Fig. 9E**). We observed DNT-like structures in contact with cells showing transfer, although major DNTs between dendrites were not imaged probably due to the weak transferred signal in the recipient dendrites (**Supplementary Fig. 9F**). This proposed transport of heavy molecules possibly shares the same mechanism with gap junction-independent transfer of calcium that we observed above (**Fig. 2K-L**).

We postulate that the DNT network is the propagation mechanism of pathological peptides in neurodegenerative brains, particularly for AD. Using our patch-infusion approach, we tracked the propagation of Human Aβ1-42 (Aβ; 4.5 kDa) by loading the Aβ peptide together with Alexa Fluor 568 (AF568; 0.7 kDa) into a single layer 5 pyramidal neuron in the mPFC (Thy1-GFP M) (**Fig. 4A**, Intraneuronal infusion; Methods). To differentiate the transfer through DNTs from the direct uptake of extracellular Aβ, an extracellular outflow experiment was conducted as the control (**Fig. 4A**). Neighboring neurons with transferred intracellular Aβ signals were observed in the infusion group but less so in the outflow group (**Fig. 4B, Supplementary Fig. 10**; Methods). The cumulative distribution of fluorescent intensities indicated the increased transfer signals by infusion (**Fig. 4C**). Specifically, the proportion of cells presenting high transfer signals (High-transfer cells; with top 5% brightness in Aβ channel) was significantly increased when the peptides were directly infused (**Fig. 4D**). This Aβ propagation was not observed in slices treated with Cyt-D, suggesting a DNT-mediated mechanism to transport intraneuronal Aβs between neurons in the brain (**Fig. 4C-D**).

**Figure 4.**
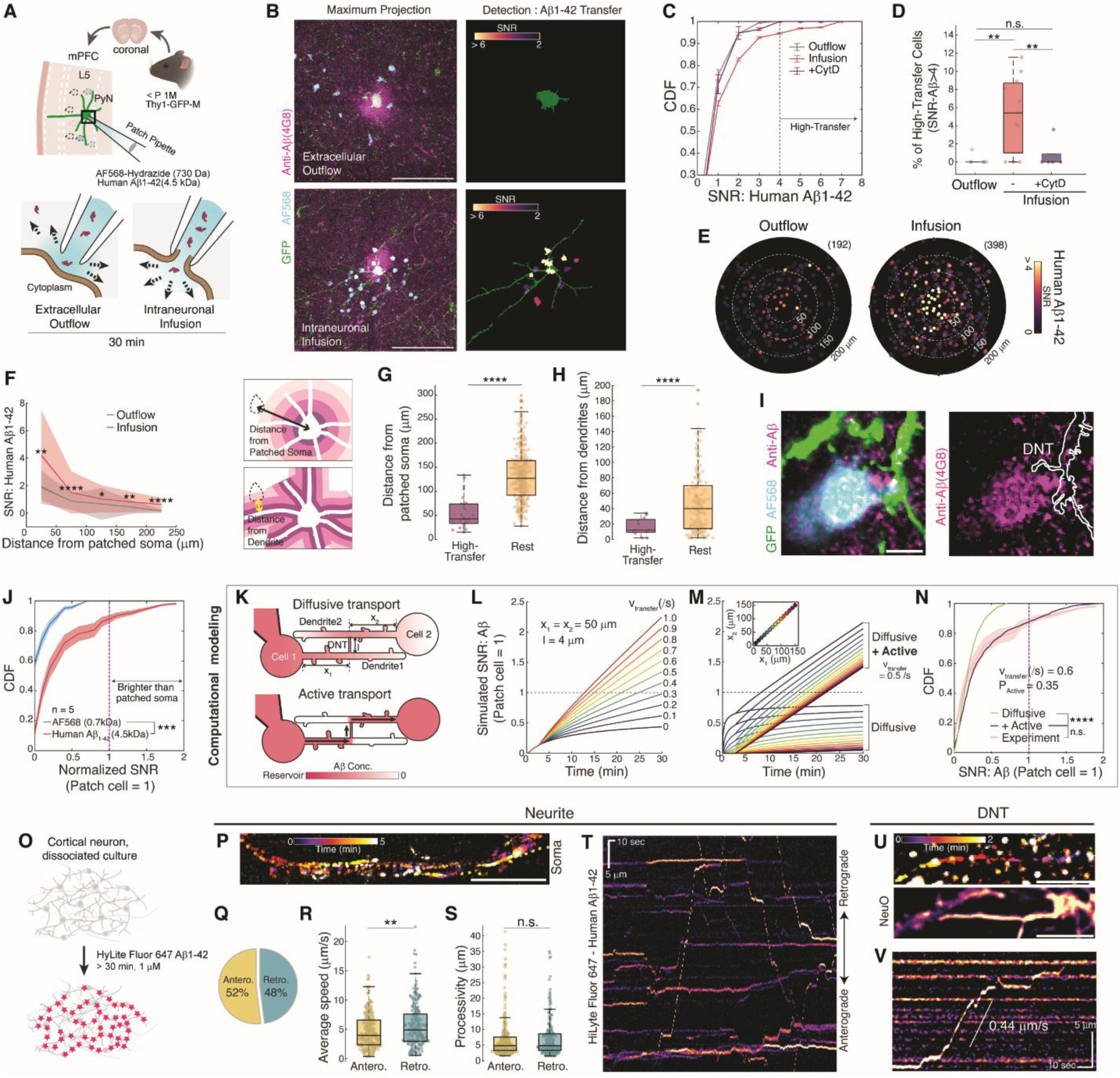
Intercellular DNT network mediates long ranged Aβ propagation. **(A)** Experimental scheme of Aβ propagation through DNT using patch clamping. The internal solution containing Alexa Fluor 568 (50 μM) and Human Aβ1-42 (1.5 μM) was either released after approaching a GFP-expressing pyramidal neuron (L5, mPFC) but not patching (extracellular outflow; bottom, left) or directly injected into the cell by whole-cell configuration (intraneuronal infusion; bottom, right) for 30 min. **(B)** Representative raw images of immunostained slices (left) for outflow (top) and infusion (bottom) groups (confocal microscopy, 40×, maximum projection of the 20 μm thick stack) and outcomes of automated Aβ transfer detection (right). **(C)** CDFs of fluorescent signals from detected recipients. To clue the mechanism of transfer, infusion experiments were also conducted in the presence of 1 μM Cyt-D in ACSF following 60-min-long pre-incubation of the brain slice (+Cyt-D). n_exp_ = 6, 11, and 5 for outflow, infusion, and infusion with Cyt-D respectively. **(D)** Fraction of high-transfer cells (SNR-Aβ1-42 > 4) from detected recipients. n_exp_ = 6, 11, and 5 for outflow, infusion, and infusion with Cyt-D respectively. *P* = 0.0021 for outflow vs. infusion, 0.0061 for infusion vs. Cyt-D, and 0.55 for outflow vs. Cyt-D by two-tailed unequal variances t-tests. **(E)** Propagation patterns of Aβ1-42 in outflow (left; n = 192) and infusion (right; n = 398) experiments. Origin = position of the paired patched soma. **(F)** Transferred fluorescent signals in recipient cells represented by their location from the patched soma. Bin width = 50 μm; line = the mean of each bin; bright shade = the standard error; dark shade = the standard deviation; two-tailed unequal variances t-test was performed to compare outflow and infusion groups for each bin. *P* = 0.0011, 0.00041, 0.022, 0.0012, and 6.5 × 10^-5^. n = 12, 53, 76, 44, and 7 for the outflow group and n = 33, 100, 144, 78, and 29 for the infusion group. **(G)** Locations of high-transfer cells measured by distance from the soma of the patched cell. *P* = 2.9 × 10^-13^ by two-tailed unequal variances t-test. n = 27 and 371 for high-transfer cells and the rest respectively, n_exp_ = 9. **(H)** Locations of high-transfer cells measured by distance from the dendrites of the patched cell. *P* = 1.4 × 10^-8^ by two-tailed unequal variances t-test. n = 12 and 166 for high-transfer cells and the rest respectively, n_exp_ = 5. **(I)** Representative high-magnification image of Aβ transfer with AF568 to a recipient cell near the dendrite of patched cell with an DNT (confocal microscopy, 63×). **(J)** CDFs of Aβ and AF568 transfer signals in recipients normalized by the signals of patched somas (n_exp_ = 5, n_cell_ = 195; *P* = 1.8 × 10^-4^ by two-sample Kolmogorov-Smirnov test; dotted line for normalized SNR = 1). **(K)** Modeling of Aβ transfer between a patched cell and a connected recipient by diffusive transport (top) and active transport (bottom). **(L)** Simulation of transfer to a recipient cell depending on the transfer efficacy (v_transfer_, the number of peptides transferred per second; v_transfer_ = 0 for diffusive transport only). **(M)** Simulation of transfer to a recipient cell depending on the path lengths (inlet, x_1_ = x_2_). **(N)** Simulated CDF for Aβ transfer in recipients (n = 2,000) when v_transfer_ = 0.6 /s and the probability of active transport (P_active_) = 0.35. *P* = 5.0 × 10^-42^ for Diffusive only vs. Active and *P* = 0.35 for Active vs. Experiment (J) by two-sample Kolmogorov-Smirnov test. **(O)** Experimental scheme to track transports of HiLyte Fluor 647 conjugated Aβs in dissociated cortical neuron (DIV11-13). **(P)** Representative time-projected image and kymograph along a neurite in time-lapse imaging (5-min-long, video rate = 1 s). **(Q)** Percentages of anterograde vs. retrograde transports (n_exp_ = 5, n_dendrite_ = 29, n_transport_ = 550). **(R-S)** Average speed and processivity of anterograde and retrograde Aβ transport. *P* = 0.0025 (R) and 0.52 (S) by two-tailed unequal variances t-test. **(T-V)** An example of time-traced Aβ transport on a DNT-like extension in dSRRF imaging of NeuO-stained live cortical neuron in time-lapse imaging (5-min-long, video rate = 0.5 s). n.s., not significant; * *P* < 0.05; ** *P* < 0.01; *** *P* < 0.001; **** *P* < 0.0001; In the box plots, the midline, box size, and whisker indicate median, 25-75^th^ percentile, and ±2.7σ, respectively. Scale bars = 50 μm (B), 10 μm (P), and 5 μm (I), (T), (U), (V).

To specify DNTs as the pathway for the transfer further, we analyzed the patterns of cellular location with the transfer, which represents long-range propagation up to > 200 μm (**Fig. 4E**). Transfer signals in recipients were observed at all distances from the patched soma (**Fig. 4G**). High-transfer cells were preferentially located near the patched soma but also near the dendrites of the patched neuron, as expected in DNT-mediated propagation (**Fig. 4H**). Moreover, DNT-like structures in contact with the recipients were occasionally observed (**Fig. 4I**), providing further evidence for the DNT network as the mechanism for Aβ propagation.

### Active Aβ transport via dendrite-DNT network

While the recipients also represented propagation of AF568, we observed differentiated transfer patterns for Aβ and AF568 (**Fig. 4E and Supplementary Fig. 6H**). The relative strength of transfer, normalized by the corresponding patched soma, clarified the distinction between the two propagations, indicating possibly distinct mechanisms for the two (**Fig. 4J**). Recipient cells never exhibited AF568 signals higher than the patched source, but their Aβ signals were higher for a portion of cells. To understand the apparent difference, we developed a computational model of cell-to-cell propagation of Aβ based on two transport mechanisms (**Fig. 4K**, Methods). The diffusive transport presumes diffusion-based propagation of peptides along dendrites and DNTs. Following this mechanism, the traveling distance along dendrites and DNTs determines the final concentration of molecules in the connected cell (Cell 2) when the concentration of the infused cell (Cell 1) is constant. Additionally, the active transport mechanism hypothesizes directional molecular transport by energy-consuming transporters, such as molecular motors, which presumably enable a consistent flow into Cell 2.

**Figure 4L** shows that for a fixed traveling distance, the Aβ concentration of the recipient (Cell 2) after 30 minutes varies greatly depending on the effective flow speed of active transport (v_transfer_). The recipient cells cannot reach the concentration (or brightness) of Cell 1 in the diffusive-transport-only model, regardless of the traveling distance, but it becomes possible when active transport coexists (**Fig. 4M**). We found parametric values to match the experimental CDF curve from 2000 simulated pairs of cells with randomly given traveling distances; 35% of DNT connections are expected to be active with v_transfer_ = 0.6 molecules/sec (**Fig. 4N**). These results suggest that AF568 propagation is diffusive-only, while specific active transport of Aβ facilitates spreading along the dendrite-DNT network.

However, no reported evidence, except active transport of APP on dendrites (*40*), supports this to our knowledge. To verify the active transport of amyloids, we tracked the dynamics of the fluorescence-tagged peptides (HiLyte Fluor 647-human Aβ1-42) internalized in dissociated cortical neurons following short (> 30min) global incubation (**Fig. 4O**; **Supplementary Movie 8**; Methods). As expected, neurites facilitated fast Aβ transport in both anterograde and retrograde directions as expected of transports by microtubule-binding kinesin or dynein motors (4.0 and 4.9 μm/s, **Fig. 4P-T, Supplementary Movie 9**). We also observed slower active transports on DNT-like interextensions, similar to the previously reported speed of TNT-associated Myosin-X (**Fig. 4U-V, Supplementary Movie 10**)(*41, 42*). Presuming most tracked signals are from aggregates, DNTs could further mediate monomers or small-sized oligomers that cannot be tracked due to photobleaching.

### Dendritic nanotube formation changes in early AD pathology

The inter-neuronal propagation of Aβ through DNTs suggests its involvement in AD pathology. We investigated Aβ-induced alterations in dye transfer via DNTs by injecting either Human Aβ1-42 or Scrambled Aβ1-42, a nontoxic peptide with a scrambled sequence of Human Aβ1-42, with a fluorescent dye AF568 (**Supplementary Fig. 9G**). No significant differences were observed in the proportion of recipient neurons (**Supplementary Fig. 9H**), the proportion of high-transfer cells (**Supplementary Fig. 9I**), or distance-based patterns (**Supplementary Fig. 9J-L**) in dye transfer from cells injected with scrambled Aβ. These results suggest that although DNTs permit Aβ transfer, they are not directly induced by Aβ at least within a short period of time.

We conducted an experiment to quantify the longer effect of global Aβ exposure on DNT formation (24-hour) in cortical neuron culture (**Fig. 5A**). When we incubate neurons in media containing Aβ, we observed amyloid accumulation inside neurons in a dose-dependent manner (**Fig. 5B-C**). We also counted the number of DNTs by using dSRRF, depending on the type of peptides (human or scrambled, **Fig. 5D**). The mild human Aβ concentration (500 nM), previously reported to increase the number of dendritic filopodia (*43*), did not significantly increase the Aβ/APP signal but did elevate the number of DNTs (**Fig. 5C-D**). Interestingly, under harsh human Aβ conditions (2 μM) (*44*), neurons exhibited a reduced number of DNTs and accumulated Aβ inside somas. These results hint the causal effect of Aβ in DNT formation, which is not linearly proportional to Aβ concentration.

**Figure 5.**
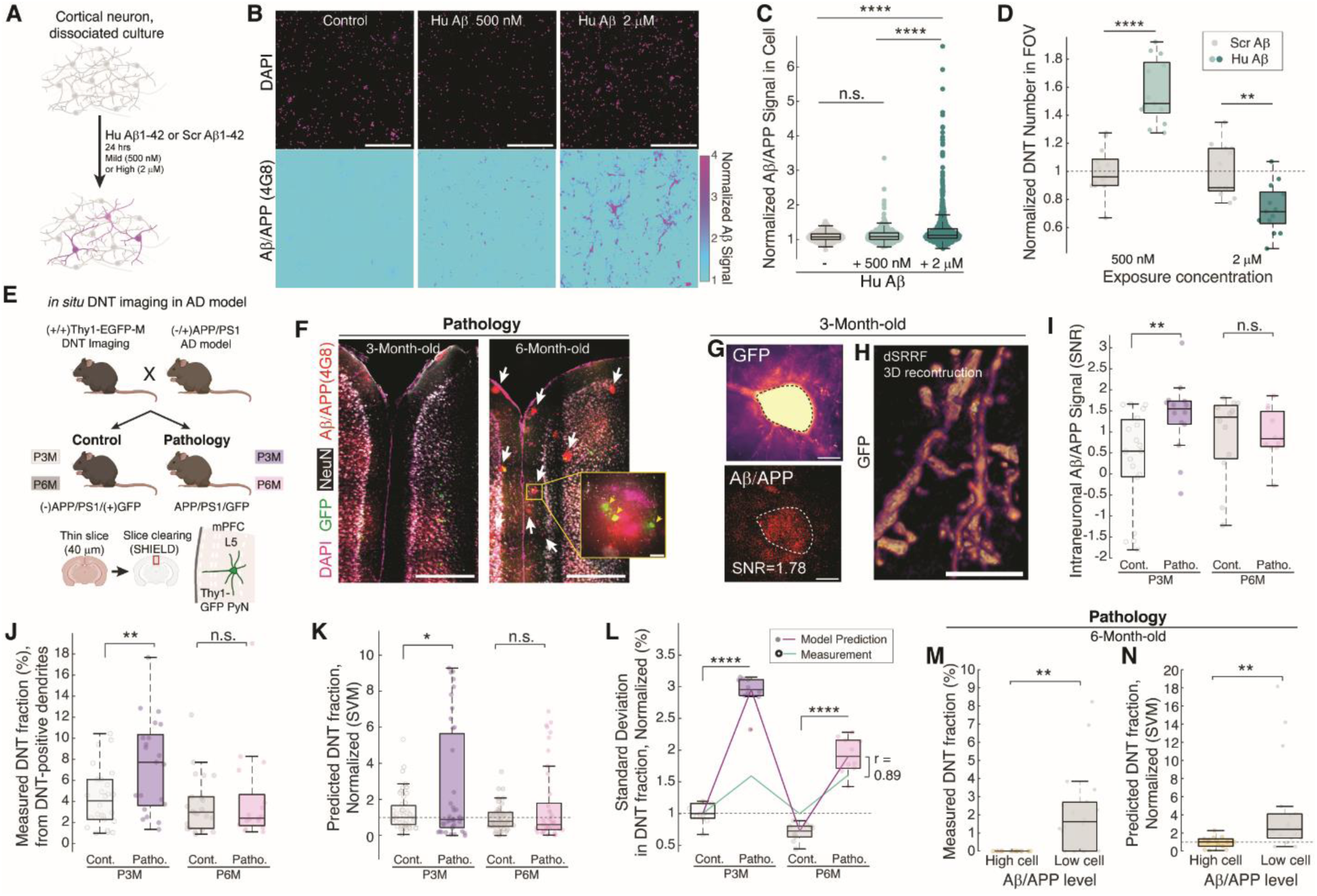
Early amyloid pathology in mouse brain exhibits altered DNT network in mPFC. **(A)** Experimental scheme of human-Aβ1-42 (Hu-Aβ) or scrambled-Aβ1-42 (Scr-Aβ) exposure on cortical neuron culture (DIV10-16). **(B)** Intracellular accumulation of Hu-Aβ in neurons after 24-hr exposure. **(C)** Normalized Aβ/APP signals in cells (n_exp_ = 3, 2, and 2; n_cell_ = 1069, 694, and 960; *P* = 0.42 for Cont. vs. 500 nM Hu-Aβ, *P* = 7.2 × 10^-35^ for Cont. vs. 2 μM Hu-Aβ, and *P* = 8.6 × 10^-22^ for 500 nM Hu-Aβ vs. 2 μM Hu-Aβ; by one-way ANOVA; *P* = 4.3 × 10^-38^). **(D)** Normalized number of DNTs in FOV (89 × 89 μm^2^; n = 13 for each condition). DNTs were defined as F-actin dominant interextensions from co-staining of Phalloidin, pNF-H, and Tau. *P* = 1.8 × 10^-5^ for 500 nM Hu-Aβ vs. Scr-Aβ, and *P* = 0.0024 for 2 μM Hu-Aβ vs. Scr-Aβ by Mann-Whitney U tests. **(E)** Experimental scheme for DNT network imaging in APP/PS1 AD model mice (n_mouse_ = 3 for each group). **(F)** Example of distinct amyloid pathology in mPFCs of P3M and P6M Pathology groups. Aβ and NeuN are detected by immunostaining (Arrows = location of amyloid plaques). (Inlet) Colocalized debris from a dead GFP-expressing neuron on an amyloid plaque (Arrows = GFP signals). **(G)** An example of intraneuronal accumulation of Aβ/APP in a GFP-expressing pyramidal neuron in the mPFC of a P3M Pathology brain. Images were maximum-projected and median-filtered for representation. **(H)** Representative dSRRF image of a DNT from a P3M Pathology brain. **(I)** Aβ signals in neurons (n = 19, 14, 12, and 12; *P* = 0.0072 for Cont. vs. Patho. in P3M; *P* = 0.89 for Cont. vs. Patho. in P6M; by two-tailed unequal variances t-test). **(J)** Measured fraction of DNTs from dendritic protrusions on DNT-positive dendrites (n = 26, 21, 22, and 17; *P* = 0.0080 for Cont. vs. Patho. in P3M; *P* = 0.74 for Cont. vs. Patho. in P6M; by two-tailed unequal variances t-test). **(K)** Mean predicted fraction of DNTs from dendritic protrusions on observed dendrites by 10 trained DNT classification models, normalized by the median of P3M Control group (n = 51, 39, 45, and 41; *P* = 0.019 for Cont. vs. Patho. in P3M; *P* = 0.068 for Cont. vs. Patho. in P6M; by two-tailed unequal variances t-test). **(L)** Standard deviation of DNT fractions in each experimental group, predicted by DNT classification models (n = 10) or measured, and normalized by the median of the P3M Control group (*P* = 8.2 × 10^-13^ for Cont. vs. Patho. in P3M; *P* = 1.6 × 10^-8^ for Cont. vs. Patho. in P6M; by two-tailed unequal variances t-test; *r* = Pearson’s correlation coefficient of two lines). **(M)** Measured DNT fractions from high Aβ/APP neurons (SNR ≥ 1; n = 15) and low Aβ/APP neurons (SNR < 1; n = 18; *P* = 0.0030 by two-tailed unequal variances t-test). **(N)** Predicted DNT fractions from high Aβ/APP neurons (SNR ≥ 1; n = 19) and low Aβ/APP neurons (SNR < 1; n = 21) by 10 DNT classification models, normalized by the median of the high Aβ neurons (*P* = 0.0089 by two-tailed unequal variances t-test). * *P* < 0.05; ** *P* < 0.01; *** *P* < 0.001; **** *P* < 0.0001; In the box plots, the midline, box size, and whisker indicate median, 25-75^th^ percentile, and ±2.7σ, respectively. Scale bars = 500 μm (B), (F), 10 μm (G), and 5 μm (H).

Elevated DNT formation by lower concentration of Aβ led us to hypothesize that DNTs play a significant role at early stages of amyloid pathology. To track the density changes of DNTs in more pathophysiological conditions, we bred Thy1-GFP-M mice and APP/PS1 AD model mice (*45, 46*), resulting in GFP-expressing neurons in the AD-pathological brain (**Fig. 5E**, Pathology; (+)APP/PS1 (+)GFP) and their controls without amyloid pathology (Control; (-)APP/PS1 (+)GFP). Given that severe amyloid pathology with senile plaques is reported in 6-month-old APP/PS1 brains (6M-Pathology), we examined 3-month-old brains as a proxy for the early pathological stage (*47*) (3M-Pathology; **Fig. 5F**). Amyloid plaques co-localized with debris of dead GFP cells were easily observed in the mPFCs of 6M-Pathology brains, supporting the concept that intracellular accumulation provides the substrates of plaque formation (*25, 26, 48, 49*) (Inlet, **Fig. 5F**). In contrast, 3M-Pathology brains did not exhibit plaque formation but showed intraneuronal Aβ/APPs (**Fig. 5G**). As expected, our super-resolution imaging successfully captured DNT structures in Pathology brains, enabling further investigation (**Fig. 5H**). We adopted a single-cell-based approach to investigate immunostained Aβ signals from the soma and morphology of dendrites protruding from each pyramidal neuron in layer 5 mPFC, where Aβ propagation was observed in previous infusion experiments.

First, we identified that 3-month-old pathology displays increased Aβ/APP signals inside neurons, corresponding to early amyloid pathology before extracellular plaques are present (**Fig. 5I**). Interestingly, intracellular accumulation was replaced by plaque conformation in 6M Pathology. To determine whether DNTs are altered in Pathology brains, we measured the fraction of manually classified DNTs on each dendrite, revealing an increased DNT fraction specifically in 3M pathology (**Fig. 5J**). Because manual quantification might not fully reflect the actual nature due to sparse fluorescent expression, we used previously trained DNT classifiers (**Supplementary Fig. 8**) to predict DNT fractions on dendrites, which reproduced the result (**Fig. 5K**). Furthermore, increases were observed in the standard deviations of DNT fractions in both manual measurements and model predictions, showing a similar trend (**Fig. 5L**). We investigated the correlation between DNT fractions on dendrites and Aβ signals inside soma. Intriguingly, we confirmed the same result from both manual and model classification; neurons with relatively high Aβ/APP signals in the soma (High Aβ Cell; **Fig. 5M-N**) particularly showed diminished DNT formation in the 6M Pathology brain. Meanwhile, amyloid-induced deformations in dendrites and axons (*50-53*) were not exhibited in our experiments, except the density of dendritic protrusion in 3M Pathology (**Supplementary Fig. 11**).

### Elevated DNT network accelerates intraneuronal accumulation in amyloid pathology

Our results verified a pathological phenotype in DNT formation during amyloid pathology in brains, particularly in the early stage (3M). However, the exact contribution of an elevated DNT network to AD pathology, particularly intracellular accumulation of Aβ, remains ambiguous. Elevated cell-to-cell Aβ propagation could either alleviate amyloid pathology by allowing cells to share the burden of degradation, as previously described in microglial TNTs (*6*), or deteriorate it by spreading the disease (*54*) as proposed previously, though it is not clearly suggested by which mechanism increased propagation systematically spreads pathology. To address how DNT networks are involved in amyloid pathology, we extended the previous computational model of active DNT-mediated transfer between two cell pairs to a lattice structure consisting of 1,668 neurons in 158 × 158 × 115 μm^3^, representing layer 5 mPFC with the measured density (**Supplementary Fig. 12A-C**; Methods). In each model, the probability distribution P_activeDNT_ for the number of DNTs with active outward transport (N_DNT_) was provided to construct the DNT network. **Figure 6A** shows three examples of representative cells whose N_DNT_s are identical to the mean DNT numbers of given P_activeDNT_ (**Fig. 6A**). Given that the proportion of neurons that displayed transfer in our Aβ infusion experiment was ∼ 13% (**Fig. 4J**) and that 35% of the total transfer was active (**Fig. 4N**), we presumed each cell to be connected to 4.6% of the neurons in the lattice on average with a 2.0% standard deviation in the normal brain used for direct infusion (P_activeDNT_ = 4.6 ± 2.0%). The probability functions were set differently to simulate pathological conditions, particularly through an increased mean and standard deviation in N_DNT_ (**Figs. 6B**, **5J-L**).

**Figure 6.**
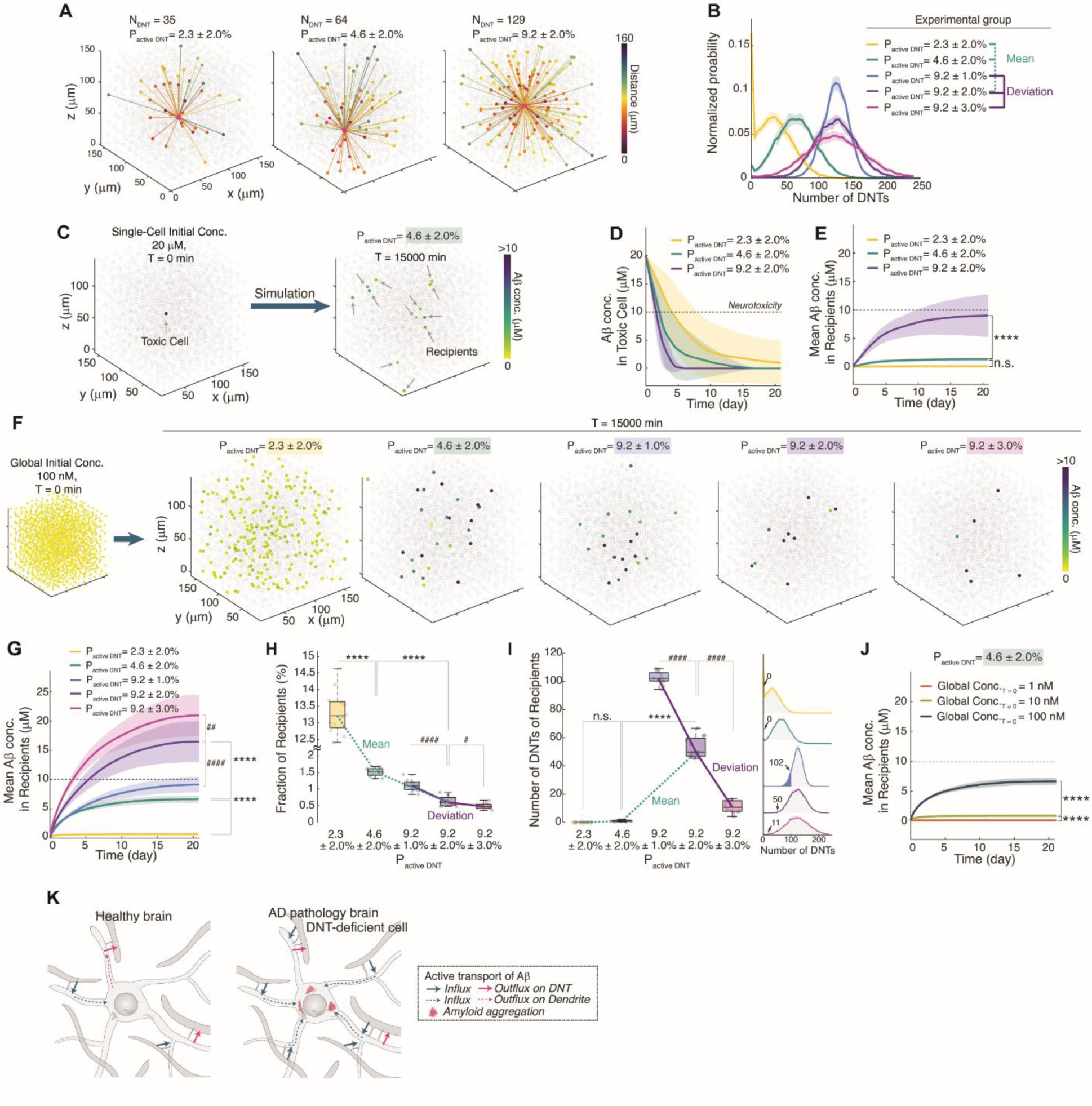
Pathological DNT network accelerates Aβ aggregation in DNT-impaired cells. **(A)** Examples of single-neuronal DNT connectivity in DNT networks simulated with different probabilities of active DNT connection between two cells (N_NB_ = the number of DNTs formed by the cell; P_active DNT_ = probability of connection by DNTs enabling active transport; colors denote the distance between connected cells). **(B)** Distributions of N_DNT_ depending on various P_active DNT_ (Comparison-by-Mean group with a fixed standard deviation = 2% and Comparison-by-Deviation group with a fixed mean = 9.2%; n = 12 for all groups). **(C)** An example of toxic cell rescue by burden-sharing in a modeled DNT network. The color of the markers represented the Aβ concentration in each cell (gray = 0; Initial Aβ concentration in a toxic cell = 20 μM where neurotoxicity in neurons appears for 10 μM). **(D-E)** Time traces of (D) Aβ concentration of the toxic and (E) mean Aβ concentration of recipient cells during burden-sharing in modeled DNT networks with different mean P_active DNT_ (n = 19 for each group). The significance test (one-way ANOVA; *P* = 1.5 × 10^-18^) was to compare the mean Aβ concentrations at the last time point (T = 15000 min; *P* = 0.25 for 2.3 ± 2.0% vs. 4.6 ± 2.0%, *P* = 5.2 × 10^-15^ for 4.6 ± 2.0% vs. 9.2 ± 2.0%). **(F)** Examples of intracellular accumulation by Aβ distribution under global amyloid exposure (100 nM) depending on different DNT network connectivity. **(G)** Time traces of mean Aβ concentration of recipient cells under global exposure in diverse simulated DNT networks with different mean and standard deviations for P_active DNT_ (n = 12 for each group). *P* = 1.3 × 10^-7^ for 2.3% vs. 4.6%, *P* = 6.5 × 10^-13^ for 4.6% vs. 9.2% by one-way ANOVA (*P* = 1.5 × 10^-18^) for the Comparison-by-Mean group. *P* = 2.3 × 10^-6^ for 1% vs. 2%, *P* = 0.0023 for 2% vs. 3% by one-way ANOVA (*P* = 1.4 × 10^-10^) for the Comparison-by-Deviation group. **(H)** The fraction of recipients for the Comparison-by-Mean group (*P* = 7.8. × 10^-38^ for 2.3% vs. 4.6%, *P* = 1.1 × 10^-5^ for 4.6% vs. 9.2% by one-way ANOVA; *P* = 4.7 × 10^-40^; P-values by the numbers of asterisks) and the Comparison-by-Deviation group (*P* = 1.4 × 10^-9^ for 1% vs. 2%, *P* = 2.0 × 10^-12^ for 1% vs. 3%, *P* = 0.039 for 2% vs. 3% by one-way ANOVA; *P* = 1.3 × 10^-12^; P-values by the numbers of hashes). n = 12 for each group. **(I)** The number of DNTs of recipients for the Comparison-by-Mean group (*P* = 1 for 2.3% vs. 4.6%, *P* = 3.9 × 10^-25^ for 4.6% vs. 9.2% by one-way ANOVA; *P* = 7.0 × 10^-27^) and the Comparison-by-Deviation group (*P* = 8.9 × 10^-21^ for 1% vs. 2%, *P* = 2.8 × 10^-29^ for 1% vs. 3%, *P* = 1.2 × 10^-18^ for 2% vs. 3% by one-way ANOVA; *P* = 6.4 × 10^-29^). n = 12 for each group. The right panel represents the location of the median N_DNT_s on the probability distributions. **(J)** Time traces of mean Aβ concentration of recipient cells during global exposure in simulated DNT networks (P_active DNT_ = 4.6 ± 2.0%) with different initial intracellular concentrations (n = 12 for each group). The significance test (one-way ANOVA; *P* = 1.7 × 10^-31^) was to compare the mean Aβ concentrations at the last time point (*P* = 1.2 × 10^-5^ for 1 nM vs. 10 nM, *P* = 8.6 × 10^-29^ for 10 nM vs. 100 nM). **(K)** Model of Aβ accumulation by DNT network in healthy (left) and AD-pathology (right) brains. * *P* < 0.05; ** *P* < 0.01; *** *P* < 0.001; **** *P* < 0.0001; In the box plots, the midline, box size, and whisker indicate median, 25-75^th^ percentile, and ±2.7σ, respectively. Bonferroni correction followed all the ANOVA tests. Line and shade are the mean and standard deviation of simulations under the same condition (D, E, G, J).

First, we validated the protective role of the DNT network by sharing the burden for degradation. A randomly selected cell in the modeled DNT network is set to have 20 μM of Aβs inside, above the concentration causing neurotoxicity (10 μM) (*55*), while the rest are amyloid-free (Single-cell toxicity). At each time interval (1 min), recipient cells connected to the original toxic cell receive Aβ influx and spread it to the next recipients. After 15,000 min (10.4 days) of simulation, we observed distributed Aβs in the network with fewer toxicities for specific final recipients (**Fig. 6C, Supplementary Movie 12**), suggesting that the burden was shared among neurons. Elevated DNT networks (P_activeDNT_ = 9.2 ± 2.0%) accelerated the speed at which the toxic cell was rescued (**Fig. 6D, Supplementary Fig.12D**). Unexpectedly, it caused severe accumulation in recipients (**Fig. 6E**). This phenomenon originates from the reduced number of decisive recipients, concentrating the burden to fewer neurons (**Supplementary Fig. 12E**).

These results suggest that selective accumulation of toxic molecules by DNT networks contributes to amyloid pathology. As we observed an increase in intracellular Aβ concentration in early AD pathology (**Fig. 5I**), we simulated progression after distributing Aβ loads evenly to all cells (100 nM in each cell) instead of setting one high-Aβ cell (Global toxicity). As expected, specific cells showed accumulated Aβs while Aβ was absent from the rest, and increased mean and deviation of P_activeDNT_ led to higher accumulations greater than the neurotoxic concentration in fewer cells (**Fig. 6F-H, Supplementary Movies 13-17**). These preferred recipients with accumulation corresponded to cells with the fewest DNTs in the model (**Fig. 6I**). An increase in the global burden developed significantly higher burdens in more specific cells (**Fig. 6J**, **Supplementary Fig. 12F-G**).

Our simulation demonstrated that the phenotype in DNT formation exhibited in our AD model can lead to intracellular accumulation, particularly in the neurons with fewer DNTs, nicely predicting the observation from 6M Pathology that high-Aβ cells represent diminished DNT formation. Thus, we conclude that DNT-mediated transport of Aβ introduces heterogeneous outcomes contextually on a tissue scale while accelerating intracellular accumulation in the context of AD progression (**Fig. 6K**).

## Discussion

Traditional studies on bridge-like structures enabling remote cell-to-cell communication have highlighted their potential significance, showcasing the diversity of cellular materials transferred along the connections. TNTs have been extensively studied due to their distinctive appearance as thin, floating nanotubular connections in cultured non-neuronal cells, which do not form any other protrusion-based connections. However, many characteristics previously attributed to TNTs do not apply to the nanotubular connectivity observed in the brain. Neurons inherently possess extensive projections and synaptic connections, so a nanotubular connection in neurons cannot be defined solely by structural connectivity over distance as for TNTs in non-neuronal cells. This would be one probable reason that we have only limited *in situ* evidences of glial TNTs, while they have been suggested to be extended to neurons *in vitro (11, 56)*.

We have adapted the definition of DNTs based on their formation mechanism, akin to cytoneme (*1, 2, 8*), which are specialized filopodia for cellular transport. Our results indicate that these actin-dominant and dynamic protrusions mediate the transfer of calcium and proteins, including amyloids. This definition includes somatic nanotubes, previously described as neuronal TNTs (*13*), presumably formed by somatic filopodia present in the early developmental stages (*14*) but diminishing as neurites mature.

DNTs exhibit distinct features from traditional TNTs. For instance, DNTs do not extend as long (∼ 3 μm; **Fig. 1K** and **3H, Supplementary Fig. 2F**) compared to TNTs in non-neuronal cells (∼ 10 μm in HeLa)(*9*). Despite this, the dendrite-DNT network facilitates distant propagation using dendrites that extend further. Additionally, we did not observe tunneling by membrane fusion at the DNT tip (**Supplementary Figs. 1, 3**), yet peptides up to 10 kDa were still transported (**Supplementary Fig. 9**). Our findings suggest connexin involvement in calcium transmission while action potential was blocked (**Fig. 2, Supplementary Fig. 7**), but gap junctions alone cannot account for the transport of larger molecules. Persisting calcium propagation after CBX treatment further supports a distinct mechanism for material transfer. Models such as transient membrane fusion, vesicular fusion, or phagocytosis might enable the transfer of large substances (*36*) without tunneling, all of which likely facilitate unidirectional transport. Unidirectional transport by DNTs formed by a donor cell to recipients is significant in DNT-involved amyloid pathology. Imbalances in input and output due to this directionality contribute to accumulation. Our computational modeling assumed a given average rate of Aβ transfer, regardless of the exact mechanism, to reflect this process.

Our evidence supports intercellular Aβ propagation via DNTs, although extracellular vesicle (EV)-mediated transfer has been suggested albeit notable limitation. In terms of quantity, the concentration of EVs in serums appears to be in the range of a few pM or lower, while the concentration of Aβ is also in the pM range in EVs released from human neural cells expressing familial Alzheimer’s disease (AD) presenilin 1 mutations (*57*) — meaning the extremely low likelihood of detecting an Aβ peptide from a million EVs. Therefore, it is less likely that the significant and specific Aβ transfer observed during infusion could be explained by the diffusion of EVs within a 30 minute window. Moreover, intracellular Aβ itself inhibits exocytosis (*58*). Interestingly, our experiments using tetanus toxin, an inhibitor of exocytosis, showed an unexpected disruption of DNT-mediated transport (**Supplementary Fig. 9M-N**), suggesting a possible shared mechanism of transient membrane fusion between EV release and DNT function.

DNTs exhibit their temporal dynamics, with rapid formation and extinction processes on a minute scale and lifetimes of a few hours (**Fig. 1L-N**). The dynamic behavior of dendritic filopodia (*32*) suggests that *in situ* DNT connectivity in the brain is plastic, responding to physiological contexts. Observation of synchronized filopodial protrusion during live-neuron imaging (**Supplementary Movie 8**) led us to hypothesize a specific signaling pathway for the temporal control of DNT formation. For instance, our recent report on the role of H-Ras in generation of nonsynaptic dendritic filopodia in pyramidal neurons suggests it as a potential regulator of DNT (*59*). This rapid response mechanism might facilitate burden-sharing through DNTs but could also accelerate amyloid pathology.

Sparse GFP expression in Thy1-GFP M model mice allows super-resolution imaging of cortical dendritic structures but reduces the likelihood of simultaneously observing binding partners of dendritic protrusions. This limitation might affect the precise measurement of DNT numbers, which is crucial for quantifying DNT-mediated connectivity. We trained classifiers with a training set of protrusions showing visible counterparts, and our Data-learners effectively distinguished DNTs from spines, estimating that DNTs constitute 12% of dendritic protrusions (**Fig. 3J, L**). However, this estimate might be exaggerated due to the binary nature of our classifiers. Additionally, some dendrites exhibited more classified DNTs (**Fig. 3L**), indicating specific dendrites contribute more to the DNT network.

In the APP/PS1 mouse model for amyloid pathology, we observed elevated DNT formation (3M Pathology) and increased deviation in the DNT fraction between dendrites (3M and 6M Pathology). This could be related to the increase in DNT-prone dendrites observed in the normal brain. Elevated cell-to-cell variance in DNT formation exacerbates input/output imbalances of amyloid at the single-cell level, especially in specific cells with deteriorated DNT connections. One possible cause for the failed DNT formation is the toxicity of accumulated Aβ which deteriorates cellular metabolism (**Fig. 1P**). Our data from amyloid-exposed cultures indicate that harsh Aβ exposure diminishes the number of DNTs (**Fig. 5D**). This suggests a pathological feedback loop where diminished DNT formation leads to more Aβ accumulation, obstructing further DNT formation. Although the change appeared statistically not significant for mild exposure, a few neurons exhibited accumulated Aβ, mirroring our simulation of global toxicity in the brain, likely trapped in this pathological loop (**Fig. 5C**). Interestingly, it was reported that treatment with Cyt-D saved dissociated neurons from amyloid-inducing toxicity (*60*), suggesting that a general decline in DNT connectivity might protect neurons from accelerated aggregation by the DNT network under global toxicity (2.3%, **Fig. 6L**).

Tracking cellular pathology in real-time is highly challenging, thus we designed computational simulations to anticipate how DNT alteration is associated in amyloid pathology. We simplified our modeling to minimize the number of input parameters and unreliable settings. Most parameters were obtained from literature research (Methods), while some key parameters were measured in our experiments. Intracellular infusion using patch clamping was a simple system to simulate, as the pipette served as a reservoir with fixed concentrations. However, real-time observation of transfer was not possible, which limited more precise modeling. Therefore, DNT connectivity was considered fixed in the modeling. Additionally, we could not exclude the effects of positive pressure flowing peptides into cells, resulting in forced diffusion in passive transfer. We also did not include the causal effect of Aβ on DNT formation because exact quantification of the effect for a given Aβ concentration was not possible. Considering this, the actual accumulation process might be delayed due to the adaptive response of forming more DNTs to increase output. Consequently, the timescale of our pathology modeling in the mPFC may be somewhat imprecise. Despite these limitations, our simulation successfully elucidates the DNT-related phenotype in the AD model brain.

Our findings highlight dendrite-DNT transport in a brain region compared to axonal transport between parts of the brain, shedding light on a novel aspect of amyloid pathology in AD (*22*). The dendrite-DNT transport network facilitates the ‘dumping’ of Aβ burden to vulnerable cells with fewer DNTs, accelerating amyloid accumulation in specific neurons. The pathological implications of intracellular amyloids in neuronal loss (*48*) or plaque formation (*49*) have been reported. DNT-mediated transport proposes a mechanism by which severe aggregation occurs even under moderate environmental concentration of amyloids by concentrating regional stress to particular cells; in our simulation, the recipients exhibited up to 200 times greater Aβ concentration than the globally given initial stress. Utilizing the DNT network, amyloid pathology may progress without leaving extracellular footprints, making amyloid-targeting treatments less efficient. It is possible that DNT-mediated intraneuronal Aβ accumulation can synergize with tau, triggering intraneuronal entanglements (*61*). Or, high intracellular concentrations might induct liquid-liquid phase separation of Aβs, accelerating fibril formation (*62*). Further studies are necessary to explore DNT-associated early pathology at the cellular level.

This study elucidated a new dimension of nonsynaptic communication between neurons in the brain. While we emphasized the pathological implications of DNT connectivity in this paper, much of its general biology remains uncharted. In contrast to well-studied synaptic interactions, DNTs can transmit not only electrical signals through calcium but also various cellular materials within a confined brain region that dendrites reach, possibly contributing to compartmentalization. Nanotubular calcium propagation facilitates distant signaling of non-neuronal cells (*5, 9, 34*), yet it remains to be discovered whether calcium transfer through DNTs is directly involved in neuronal activity. Moreover, DNT connections might extend to neuron-glia interactions in the brain, as glial nanotubular structures have been reported (*6, 10, 56*). Our research highlights the need for further investigation to reveal how DNT connectivity shapes communication among different types of brain cells (**Supplementary Fig.6G**), including the urgent need to develop DNT-specific markers for easier access to these structures.

## Acknowledgments

We thank members of the Kwon laboratory and Johns Hopkins Neuroscience community; C. Kwak, K.Nagahama, S.Roh, M.Xiao, J. Troncoso, and P.Worley for helpful discussion. APP/PS1 mice were generously provided by R.Huganir. We thank A.Bush of Neuroscience Imaging Core and Y.K.T.Xu of D.Bergles laboratory for helping with brain clearing and light sheet imaging. We thank M.Chang, P.Kanold, and S.Blackshaw for sharing the ephys setup and cell culture room.

## Author contributions

M.C. and H-B.K. designed this study and wrote the manuscript. S.K. performed Aβ infusion experiments with patch clamping. L.K.P. inspected DNT formation in FIM/SEM images of mouse brains. J.Kim. performed dye infusion experiments with patch clamping. D.L., A.M., J.Kwon. helped mark DNTs from images, prepare samples, and develop modules for data import. S.O. provided advice on the manuscript. All other tool development, experiments, analysis, data interpretation, and computational simulation were performed by M.C.

## Methods

### Animals

Thy1-GFP line M mice breeding pairs (Cat.# 007788) from the Jackson Laboratory (Bar Harbor, ME, USA) were purchased and inbred to generate the homozygous genotype used in this study. Thy1-GFP-M mice were used for morphological imaging (1-8 months old, both sexes; n_mouse_ = 5 in total) and whole-cell patch-clamp infusion (2-4 weeks old, both sexes; n_mouse_ = 24 in total). APP/PS1 AD model mice were generously provided by Richard Huganir Lab (Johns Hopkins University School of Medicine, Baltimore, MD, USA) and bred with homozygous Thy1-GFP-M to obtain both APP/PS1(+)/GFP(+) and APP/PS1(-)/GFP(+) mice, which were used for AD study (3 and 6 months old; n_mouse_ = 12 in total). Both female and male animals were randomly selected for experimental groups. For the dissociated culture of cortical neurons, timed-pregnant (E15) mice were purchased from the Jackson Laboratory. Genotyping of mice used in this study was routinely performed when the animals reached 3 weeks old (TransnetYX, Cordova, TN, USA). All procedures were carried out under protocols approved by the Johns Hopkins University Animal Care and Use Committee (protocol number MO22M170) and the National Institutes of Health guidelines.

### Preparation of dissociated cortical culture

Dissociated culture of cortical neurons was carried out as previously described but slightly modified to mice (*63*). On the next day of arrival (E16), pregnant mice were euthanized and cortices of embryos were dissected. Collected cortices were incubated in 0.67 mg/ml of papain (Worthington, Lakewood, NJ, USA, Cat.#: LS003119) and 0.001% DNase (Sigma-Aldrich, St.Louis, MO, USA, Cat.#: DN-25) for 15 min at 37°C. Cells were carefully triturated with a glass pipet and 1,000 μL-sized pipet tip ten times, respectively. Dissociated neurons were cultured in 24 well glass bottom plates (Cellvis, Mountain View, CA, USA, Cat.#: P24-1.5H-N) or 4 well chambered cover glass (Cellvis, Cat.#: C4-1.5H-N) which were PDL-coated by 2-hour-long incubation of 40 µg/ml PDL (Sigma-Aldrich, Cat.#: P0899) at RT. The final density of cells was 1 × 10^5^ per well. For time-lapse imaging, the density was reduced to 2.5 - 5 × 10^4^. Neurobasal medium (Thermo Fisher Scientific, Waltham, MA, USA, Cat.#: 21103049) was used for culture supplemented with 5% (v/v) HS (Thermo Fisher Scientific, Cat.#: 26050070) or FBS (Cytiva, Marlborough, MA, USA, Cat.#: SH30071.02), 1% (v/v) Glutamax Supplement (Thermo Fisher Scientific, 35050061), 2% (v/v) B27 supplement (Thermo Fisher Scientific, Cat.#: 17504044), and 1% (v/v) penicillin-streptomycin(Thermo Fisher Scientific, Cat.#: 15070063). Primary cortical neurons were grown at 37°C and 5% CO_2_ conditions. Every 3∼4 d, a half volume of media was replaced with fresh media with 1% serum. 2.5 μM Ara-C (Sigma-Aldrich, Cat.#: C-6645) was added for the first media replacement to prevent glial growth.

### Sample preparation for SRRF imaging of cultured cortical neuron

For SRRF imaging, cortical neuronal culture (DIV7 or DIV14) was fixed as previously described (*31*). Initially, cells were fixed for 1–2 minutes using a solution containing 0.3% (v/v) glutaraldehyde and 0.25% (v/v) Triton X-100 in cytoskeleton buffer (CB, consisting of 10 mM MES, pH 6.1, 150 mM NaCl, 5 mM EGTA, 5 mM glucose, and 5 mM MgCl_2_). Subsequently, they were post-fixed for 10 minutes in 2% (v/v) glutaraldehyde in CB. Finally, the sample was treated with freshly prepared 0.1% (w/v) sodium borohydride for 7 minutes to diminish background fluorescence. For DNT modulating experiments, culture media were exchanged to expose cells to 0.1% DMSO or desired drug (1 μM Cytochalasin-D, Sigma-Aldrich, Cat.#: C2618) for 90 minutes before fixation.

Fixed cells were incubated for > 1 hr in blocking buffer (3% (w/v) BSA, 0.2% (v/v) Triton X-100 in PBS) for the further staining procedure. First, immunostaining of neurofilament (1:200 dilution, AF488 anti-NF-H, phosphorylated antibody, BioLegend, San Diego, CA, USA, Cat.#: 801612) and tau (1:200 dilution, AF647 Tau-46 antibody, Santa Cruz Biotech, Dallas, TX, USA, Cat.#: sc-32274) was carried out for overnight at 4°C. After washing, F-actin staining with 500 nM AF568 conjugated Phalloidin (Thermo Fisher Scientific, Cat.#: A12380) followed for 1 hr at RT. 300 nM DAPI was added in the final step for nucleus staining.

### Tissue fixation, clearing, and sample preparation for brain slice SRRF imaging

For tissue fixation animals were deeply anesthetized by isofluorane and then perfused transcardially with PBS (pH 7.4) and with 4% paraformaldehyde (PFA). The brains were collected and postfixed in 4% PFA overnight at 4°C. A vibratome (Leica VT 1000S) was used to slice coronal sections (40 μm) from the brain region of interest (**Supplementary Fig. 3**). Before tissue clearing, brain slices with optimized GFP expression for imaging were selected by checking it with the low magnification objective (10×). Modified SHIELD protocol was conducted for slice clearing (Lifecanvas, Cambridge, MA, USA) (*64*). Briefly, PFA-fixed brain slices were incubated in the mixture of SHIELD Epoxy and SHIELD ON solutions (1:7 ratio) for > 6 hours at 4°C, then kept shaking at RT overnight. The next day, slices were washed with PBS and put in Passive Clearing Buffer for 6 hours at RT. Cleared tissues were washed in 0.05% PBST at 37°C overnight. For immunostaining of intracellular Aβ in AD-model studies, primary antibody incubation (4°C, overnight; BioLegend, Cat.#: 800710) and secondary staining (37°C, 2 hours; Thermo Fisher Scientific, Cat.#: A32728) followed after 6-hour-long storage in blocking buffer (1% BSA, 0.3% Triton X-100 in PBS). Prepared brain slices were mounted on a microscope slide with EasyIndex (Lifecanvas) and sandwiched with a precision coverglass (#1.5H, Thorlabs, Cat.#: CH15CH or CH15KH).

### Microscopy

Most of images used in this study were obtained by a home-built microscope specifically optimized for SRRF imaging and live-cell imaging of cultured cells and cleared thin brain slices (**Supplementary Fig. 3A**). Briefly explaining, a manually aligned highly-inclined illumination system of lasers (CellX Laser 4 x 100 mW 405:488:561:637 nm, Coherent, Saxonburg, PA, USA) and an extra LED illumination system (X-Cite XYLIS, Excelitas, Waltham, MA, USA) combined into an inverted microscope (IX83 2-deck system with z-drift compensator (IX83-ZDC2), Evident Scientific, Waltham, MA, USA) enabled high-resolution imaging by high-magnification objectives (U Plan S-APO 60× silicone NA 1.3, WD 0.3 mm, Evident Scientific) and wide field-of-view imaging (U Plan S-APO 30× silicone NA 1.05, WD 0.8 mm and U Plan FL 10X Phase, PH1, NA 0.3, WD 10mm, both from Evident Scientific) respectively. A customized laser quad band filter set (ZT405/488/561/647rpc & ZET405/488/561/647m, Chroma, Bellows Falls, VT, USA) was installed to the highly-inclined illumination system, which facilitated multi-color imaging by fast laser-alteration without filter alteration. iXon Life 888 EMCCD (1024 × 1024, pixel size = 13 μm) was used for the detection of emitted fluorescent signals while a customized lens system (Thorlabs, Newton, NJ, USA) matched the final actual pixel size to be ∼ 100 nm when imaged by the 60× objectives. All optical components were connected and controlled by Metamorph software advanced for Olympus optical system (Molecular Devices, San Jose, CA, USA). Home-built journal scripts and Multi-Dimensional Assay application for Metamorph empowered all necessary operations. Focal position and actual pixel size were measured using an Invitrogen FocalCheck Fluorescent Microsphere Kit (6 μm, mounted on slides, Thermo Fisher Scientific, Cat.#: F24633).

For the single-cell uncaging experiment, an additional illumination module including 365 nm LED light (190 mW, 700 mA, Thorlabs) was incorporated through a dichroic beamsplitter (376 nm, Semrock, Rochester, NY, USA) in the quad-laser highly-inclined illumination path. A 4f relay lenses configuration with a precision pinhole (50 μm, Thorlabs) shaped point focus on the focal plane of other illuminating lasers. The fixed point focus was measured from the images of 7-Diethylamino-4-methylcoumarin (100mg/ml in distilled water; Sigma-Aldrich, Cat.#: D87759) to be a diameter of ∼ 20 μm (Otsu’s thresholding; ∼16 μm in FWHM). A ½’’ mechanical beam shutter and *Kinesis* software (Thorlabs) enabled temporal control of UV exposure.

### Whole brain tissue clearing and light sheet imaging

The brain of a 1-month-old Thy1-GFP M mouse was prepared following CUBIC L Tissue Clearing (*37*) and refractive index matching (RIMS)(*38*) protocols that were optimized for brain imaging. Cleared brain was imaged by 3D light-sheet imaging (Zeiss Lightsheet 7; magnification of detection objectives = 20X with 25X zoom, voxel size = 0.184 × 0.184 × 1.116 μm^3^). Sample preparation and imaging protocols were provided from Dwight Bergles Lab and the Neuroscience Imaging Center (Johns Hopkins University).

### dSRRF image acquisition

Deconvolution-assisted SRRF (dSRRF) was performed for volumetric reconstruction of neuronal structures of brain slices and dissociated culture in super-resolution. For cultured neurons, 100 - 1000 frames of images were acquired with an acquisition rate of 30 fps (FOV = 650 × 650 pixel^2^ or 69×69 μm^2^). For multi-color reconstruction, image stacks for each fluorescent signal were sequentially obtained by alternating the wavelength of the excitation laser. The intensities of the excitation lasers were set so that the fluorescent signal in the field of view was maximized without saturation when the EM gain of the EMCCD was set to 3000. NanoJ-SRRF (*65*), the imageJ plugin for GPU-assisted SRRF analysis (NVIDIA GeForce RTX 2060 SUPER), executed the reconstruction process (parameter setting of ring radius = 0.5, radiality magnification = 3, axes in ring = 6, temporal radiality pairwise product mean, renormalize, with intensity weighting).

For every given axial displacement of 500 nm, an image sequence of 100 frames was obtained to scan prepared brain slice samples (40 μm) following tissue clearing. A single volumetric image stack for reference was obtained beforehand. The acquired set of image sequences was decomposed and recombined to make 100 image stacks of the volume. GPU (NVIDIA GeForce GTX 1080 Ti)-assisted fast deconvolution of the image stacks by the Richardson-Lucy algorithm was executed by a Matlab script where the point spread functions for each fluorescence were generated by PSF Generator, an imageJ plugin (Biomedical Imaging Group at EPFL), and used for the procedures. Deconvoluted stacks are recombined into image sequences, reconstructed by NanoJ-SRRF, and drift-compensated (Matlab) to yield a super-resolved volumetric stack. See Supplementary Figure 2 for more information.

### Characterization of DNT from H01 dataset

The EM-resolved morphology of dendritic protrusions in the human brain was investigated using Neuroglancer (https://h01-dot-neuroglancer-demo.appspot.com/). Dendritic filopodia or spines with clear heads were randomly selected and classified as nonsynaptic or synaptic by the provided synapse marking in Neuroglancer. For identification of nonsynaptic filopodia, serial axial scanning followed for confirmation. To investigate contact to other dendrites from protrusions, we checked all the axial planes containing the terminal of the protrusion and identified all the contacting segments. End-to-end distance and thickness of head were measured as the distances between two coordinates of the tip and the bottom and the farthest points in an axial plane showing the thickest head. Movies of serial axial planes were made from copies of axial planes using ImageJ.

### Characterization of DNT from SRRF images of dissociated neurons

Reconstructed dSRRF images were analyzed using ImageJ and Matlab. interextensions between neurites were manually marked using Freehand lines and saved as ROIs (ROI Manager, ImageJ). A neurite was defined as a projection from a soma or a branch from other neurites. Some neurites were saved for comparison with interextensions. Saved ROIs and corresponding multicolor SRRF images were loaded using ReadImageJROI (Dylan Muir, 2014) and loadtiff (YoonOh Tak, 2012) functions on Matlab and analyzed further. For each freehand line ROI, the length of the structure was measured as the end-to-end distance. To measure the local width and SNR, we obtained intensity profiles from a hypothetical line perpendicular to the ROI (*l* = 2 μm) for each pixel point on the ROI line. These profiles were walking-averaged by 5 pixels and used for local FWHM (full-width-half-maximum) measurement. At the same time, local SNR was measured from the peak value of the same hypothetical line on the original image before SRRF processing. To decide SNR, the mean and noise of the background were defined as the mean and standard deviation of gray values in the local background, which was marked by binarizing bounding region (10 μm × 10 μm) with *adaptthresh* (sensitivity = 0.6, neighborhood size = 1.6 μm). The width and SNR of the extension were defined as the minimum local width and the mean local SNR.

### Time-lapse tracking of DNTs in dissociated neuronal culture and analysis

Cortical neurons in reduced density culture were used for the time-lapse experiment when they reached DIV 1-7 because it became difficult to distinguish single filopodium in older cultures with heavy projections. Our home-built fluorescent microscope equipped with a Stage Top Incubator (Tokai Hit USA Inc.,Bala Cynwyd, PA, USA) is used for imaging while maintaining temperature (37°C) and CO_2_ concentration (5%). The objective lens heater was also used and the correction collar of the objective was set for 37°C. For time-lapse imaging, the culture medium was exchanged with BrainPhys Imaging Optimized Medium (Cat.#: 5796) supplemented with NeruoCult SM1 (Cat.#: 5711) and 200nM NeuroFluor^TM^ NeuO (Cat.#: 1801; all from STEMCELL Technologies, Vancouver, BC, Canada) to enable filopodial imaging of neurons while minimizing background fluorescence from phenol red. While 1-hr-long pre-incubation, cells were transferred to the stage top incubator for thermal equilibrium which is essential to minimize thermal drift during imaging. For each 1 – 2 hrs imaging session, multiple locations (10 – 12) were imaged every 30 or 60 sec to increase the chance of observation.

Whenever the identical location was imaged, axial drift compensation was performed. A 561 nm laser was used for the excitation of NeuO-stained protrusions. Acquired image stacks for each location were analyzed using ImageJ to mark DNTs and count the number of frames in which those structures appeared. To test the pharmacological effects on DNT formation, 0.1% DMSO, 2 μM Rotenone (Sigma-Aldrich, Cat.#: R8875), 1 mM ATP (Roche, Basel, Switzerland, Cat.#: 11140965001), and 1 μM Cytochalasin-D (Sigma-Aldrich, Cat.#: C2618) were tested in a randomized order from wells of the same cultures. For temporal stability measurement of filopodia-forming extensions, DIV1 neurons were stained with 100 nM SiR-actin (Cytoskeleton, Inc., Denver, CO, USA, Cat.#: CY-SC001) for 66 hours (video rate = 15 min). Imaging was performed as previously described.

### Immunohistochemistry in dissociated neuronal culture

Cortical neurons cultured in dissociation (DIV16) were fixed with 4% PFA for 30 minutes, washed, and immunostained with Rabbit Anti-CaMKII (Sigma-Aldrich, Cat.#: C6974), Mouse Anti-Parvalbumin (Swant, Burgdorf, Switzerland, Cat.#: PV235), and Rat Anti-Somatostatin (Sigma-Aldrich, Cat.#: MAB354; 1:300 for all). After incubation for overnight, 1-hr-long secondary staining followed with Goat Anti-Mouse IgG-AF488 (Thermo Fisher Scientific, Cat.#: A11029), Goat Anti-Rabbit IgG-AF568 (Thermo Fisher Scientific, Cat.#: A11036), Goat Anti-Rat IgG-AF647 (Thermo Fisher Scientific, Cat.#: A21247; 1:300 for all antibodies), and 300 nM DAPI. Images were analyzed by segmenting cells from the DAPI channel (adaptive thresholding), measuring each immunostained signal in signal-to-noise ratio, and comparing the SNRs to decide the brightest signal using Matlab.

### Single-cell calcium uncaging and transfer

Cortical neurons cultured in dissociation and grown for 13 – 17 days were used for the experiment. Cells were moved to the stage-top incubator of the microscope which had been prepared to achieve and sustain the preset temperature (37°C) and CO_2_ concentration (5%) beforehand (> 60 minutes). Afterward, cells were incubated with 20 μM DMNPE-4-AM(R&D Systems, Minneapolis, MN, USA, Cat.#: 5948/5), 5 μM Cal-590-AM (AAT Bioquest, Pleasanton, CA, USA, Cat.#: 20512), 10 μM Verapamil (Sigma-Aldrich, Cat.#: 676777), 200 nM NeuO, 0.1% Pluronic F-127 (Sigma-Aldrich, Cat.#: P2443), 2.5 mM Probenecid (AAT Bioquest, Cat.#: 20061) in BrainPhys Imaging Optimized Medium supplemented with NeruoCult SM1. Verapamil, a calcium channel blocker, was added to inhibit calcium entrance through channels and action potential firing during the experiment (*66*). Depending on the experiment, 0.1% DMSO (Thermo Fisher Scientific), 1 μM Cytochalasin-D (Sigma-Aldrich, Cat.#: C2618), 100 μM Carbenoxolone (Sigma-Aldrich, Cat.#: C4790), 50 μM BAPTA-AM (Tocris, Minneapolis, MN, USA, Cat.#: HY-100545), 100 μM D-AP5 (Tocris, Cat.#: 0106), 20 μM Nimodipine (Tocris, Cat.#: 0600), 30 μM Trovafloxacin (Tocris, Cat.#: 3863) was added with or without 1 μM Tetrodotoxin citrate (Tocris, Cat.#: 1069) while the amount of DMSO for the control group was set to match the amount of Cyt-D or CBX. These were tested in a randomized order from wells of the same cultures. Following 60 – 90 minutes long pre-incubation, single-cell uncaging experiments were performed. For each trial, the locations of cells were imaged using 488 nm which excites NeuO (FOV = 1024 × 1024 pixel^2^ or 94 × 94 μm^2^). The position of a *target* cell was located on the pre-checked UV point focus. After acquiring the image as a reference, the excitation laser was switched to 561 nm for Cal-590-AM indicator imaging. An image sequence was obtained for 100 sec including the pre-exposure session (30 sec), UV-exposure session (50 sec), and post-exposure session (20 sec) with a frame rate of 10 Hz (1,000 frames in total). UV exposure was controlled by opening the bean shutter for 50 sec following a user input on the 30 sec. For each well, 15 – 20 target cells were selected for calcium uncaging and transfer imaging. For each experimental group, data was collected from 2 wells.

### Analysis of calcium transfer

Collected image sequences were resized to 512 × 512 pixel^2^ for analytic convenience. First, the boundaries of somas for *targets* and *neighboring cells* were manually marked by blind researchers using Freehand selections and saved as ROIs (ROI Manager, ImageJ). Home-built Matlab scripts performed further analysis. Saved ROIs and corresponding image sequences were loaded using ReadImageJROI (Dylan Muir, 2014) and loadtiff (YoonOh Tak, 2012) functions. Time-dependent mean gray values and center-of-mass of each ROI were saved as a time trace and the location of individual cells. For each experiment, the initial and final time points of UV exposure were specified from the trace of the *target* by checking stepwise increase and decrease of intensities due to direct UV detection from our EMCCD. Afterward, the traces from *neighbors* in the same image sequence were inspected to exclude traces showing synchronized stepwise increase and decrease above noise level, which indicate direct exposure of UV to the cells. These filtering processes successfully excluded traces with spontaneous firing which are not synchronized with UV on-off timing. All the traces were grouped by their experimental condition and analyzed together. For the representation of traces, traces were realigned to match the initial time points of UV exposure.

#### Generation of surface plot

For each time trace of calcium indicator intensity in *neighboring* neurons, the time-dependent representatives were obtained by averaging the trace values for the time intervals of 10-20 sec, 30-40 sec, 50-60 sec, 70-80 sec, and 90-100 sec. These time-representatives were classified by the distance between the *neighbor* and *target* into the bins of 20-30 μm, 30-40 μm, 40-50 μm, and 50-60 μm. For these time-distance bins, we calculated the ratio of the number of cells with ΔF/F_0_ > 0.1 as a proxy of the transfer probability. The resultant 3-dimensional array (5 X 4) was represented using the *surf* function followed by *shading* with interpolation on Matlab. A user-developed colormap is used for representation (YlGnBu, *slanCM*; Zhaoxu Liu, 2024.).

### Morphological analysis of *in situ* DNT

Reconstructed dSRRF images of GFP neuronal morphology were analyzed using ImageJ and Matlab in a similar method as described above for dissociated neuron culture but modified for 3-dimensional structures. Dendritic protrusions were marked using Freehand lines and saved as ROIs in ROI Manager (ImageJ) by blind researchers. The line was drawn to cover the whole protrusion in a maximum-projection image. Saved ROIs and corresponding multicolour SRRF images were loaded using ReadImageJROI (Dylan Muir, 2014) and loadtiff (YoonOh Tak, 2012) functions on Matlab and further analyzed. Morphological characterization of protrusions was conducted on the axially-averaged projection image (5 images from 2 μm thick sections). Local intensity profiles were obtained from hypothetical lines perpendicular to each ROI (*l* = 40 pixels) for pixel points on the ROI line. These profiles were walking-averaged by 3 pixels and used for local FWHM (full-width-half-maximum) measurement, yielding local widths. Local peak intensities were measured from the peak value of the same profiles and converted into SNR (peak intensity from mean background gray values divided by the standard deviation of background gray values) and ΔF/F_0_ (peak intensity from mean background gray values divided by the mean background gray values). Mean and median absolute deviations of the local widths and peak intensities were also measured. Structural similarity index (SSIM) was obtained using *ssim* funciton (Matlab). Curve Fitter App from Curve Fitting Toolbox of Matlab was used for Gaussian fitting of probability distributions.

#### DNT classification learner

Quadratic support vector machine (SVM) models were trained with 22 morphological features (path length, end-to-end length, maximum, minimum, mean, and median peak intensities in SNR and ΔF/F_0_, mean and median absolute deviation in peak intensities, skewness for x and y, kurtosis for x and y, maximum, minimum, mean, and median of widths, mean and median absolute deviation in widths) of DNTs (n = 83) or axon-bound dendritic spines (n = 176) to generate trained classifiers (**Supplementary Fig. 4B**). Each classifier was trained with 80% of the total data set that is randomly partitioned and tested with the remaining 20% to validate its decoder performance by the area under the ROC curve (AUC; **Supplementary Fig. 4C**). To train Shuffle learners, the labels of the actual data were randomly permutated and reassigned to generate the shuffled data set. Classification Learner App provided in Machine Learning Toolbox of Matlab was used for training and analysis.

### Preparation of acute cortical slices

Acute slices containing the mPFC were prepared from 2–4-week-old Thy-GFP-M mice in aseptic conditions. Mice were deeply anesthetized with isoflurane and cardiac perfused with ice-cold artificial cerebrospinal fluid (ACSF) solution (in mM: 124 NaCl, 3 KCl, 1.3 MgSO_4_, 2.5 CaCl_2_, 10 D-Glucose, 1.25 NaH_2_PO_4_, NaHCO_3_) prior to decapitation. The brains were quickly removed and submerged in ice-cold cutting solution (in mM: 215 Sucrose, 20 D-Glucose, 26 NaHCO_3_, 4 MgSO_4_, 4 MgCl, 1.6 NaH_2_PO_4_, 1 CaCl_2_, 2.5 KCl). Cortical slices (300 μm thick) were prepared using a VT1000S vibrating microtome (Leica) and then incubated at 30°C ± 2°C for 30 minutes in a holding chamber filled with ACSF. The holding chamber was moved to RT and slices recovered for 30 more minutes before recording. All solutions were saturated with 5% CO_2_/95% O_2_ during and at least 30 min prior to slice preparation.

### Patch clamping and molecular infusion

Acute brain slices were transferred to a submersion-type recording chamber perfused with oxygenated ACSF (in mM: 124 NaCl, 3 KCl, 1.3 MgSO_4_, 2.5 CaCl_2_, 10 D-Glucose, 1.25 NaH_2_PO_4_, NaHCO_3_). Whole-cell recordings were performed at room temperature on visually identified and transfected layer 5 pyramidal neurons of the mPFC using a MultiClamp 700B amplifier (Molecular Devices) and an upright microscope (E600 FN, Nikon) with oblique infrared and fluorescent illumination. Neurons were patched in voltage-clamp configuration (Vhold = -65 mV) using borosilicate glass pipettes (electrode resistances 3-7 MΩ) filled with internal solution (in mM: 2.7 KCl, 120 KMeSO_4_, 9 HEPES, 0.18 EGTA, 4 MgATP, 0.3 NaGTP, 20 phosphocreatine (Na), pH 7.3, 295mOsm) which conditionally included Alexa Fluor 568 Hydrazide (50 μM, Thermo Fisher Scientific, Cat.#: A10437) together with either human Aβ1-42 (1.5 μM, Anaspec, Fremont, CA, USA, Cat.#: A24224), scrambled Aβ1-42 (1.5 μM, Anaspec, Cat.#: A25382), or Dextran-Alexa Fluor™ 488 (50 μM, Thermo Fisher Scientific, Cat.#: D22910, MW = 10 kDa). Data was acquired with the “WaveSurfer” plugin tool in MatLab. To characterize cell types of the recorded cells, we switched to current-clamp and injected -100 – 500 pA depolarization current steps to evoke action potentials (50 pA increase per step). Only neurons with a stable steady-state holding current (<200 pA) were considered for further analysis. To allow for complete internal solution infusion and propagation along dendrite-DNT network, cells were kept in a whole-cell patch for 30 minutes. High-amplitude, high frequency depolarization current steps (10 nA at 100 Hz for 100 ms) were injected into the recorded cells at the end of recording to increase efficiency of intracellular infusion as previously described (*67, 68*). As control, we allowed the flow of internal solution into the extracellular space of our targeted, GFP-expressing neuron without initiating a whole-cell patch, ensuring that transfer was initiated via DNTs and not exo- and endocytosis. Slices were immediately fixed after recording by dropping 4% PFA solution on them and prepared for further analyses.

For the DNT-disruption experiment, slices were incubated for 60 minutes before transfer to the recording chamber and perfused during recording with the ACSF saturated with 5% CO_2_/95% O_2_ with 1 μM Cytochalasin-D (Sigma-Aldrich, Cat.#: C2618). For the experiment to test the effect of tetanus toxin in Aβ transfer, 100 nM tetanus toxin (Sigma-Aldrich, Cat.#: 582243) was added to the internal solution.

### Immunohistochemistry and fluorescence imaging of Aβ transfer in acute slices

Fixed slices were washed with PBS, incubated for permeabilization (1% BSA, 1% Triton X-100 in PBS; RT, 1 hr), and incubated for 48 hrs for primary immunostaining with Chicken anti-GFP (1:400; Abcam, Cat.#: ab13970) and Mouse anti-Aβ (1:200; clone 4G8, BioLegend, Cat.#: 800710) at 4°C. Afterward, secondary immunostaining followed at 37°C for 2 hrs (Goat anti-Chicken IgY-AF488, Thermo Fisher Scientific, Cat.#: A11039; Goat anti-Mouse IgG-AF647, Thermo Fisher Scientific, Cat.#: A32728). The stained slices were mounted on the slide glass with a mounting solution (DAPI Fluoromount-G, SouthernBiotech, USA) and imaged using Zeiss LSM 880 confocal microscopy to obtain 4-colored, tiled image stacks (DAPI / GFP-AF488 / AF568 / Aβ-AF647; lateral pixel size = 297 nm and axial step size = 2 μm with 40× objective; Neuroscience Imaging Center, Johns Hopkins University). The approximate location of the patched cell was recovered using the image recorded during patch clamping as a reference. The precise patch cell was specified by the image in the GFP channel, which was previously recorded during patch clamping and had its background signal (AF568) increased locally around the cell. Occasionally we observed a locally increased background but no patch cell itself. These images were considered as from slices with dislocated soma after infusion, and only used for transferred signal measurements for recipients (**Fig. 4C-E**) but not for distance-based analysis. To measure the distance of recipients from GFP dendrites, the best experimental images that enable dendrite tracking were selected (n = 5).

We found that clone 4G8 is more efficient for staining of intracellular Aβ than clone 6E10, though the antibody is less ideal in terms of its specificity which detects both human Aβ and mouse APP (**Supplementary Fig. 10**). For the high concentration of human Aβ was infused in this experiment, we regarded detected Aβ signals as a measure of amyloid transport.

### Analysis of DNT-mediated Aβ transfer in acute slices

Home-built Matlab scripts were used for the automated quantification of transferred signals in recipient cells. The images containing blood vessels with strong staining were excluded from the analysis. For images to be analyzed, the axial range of interest was set from each stack to include the patched cell but exclude a couple of frames from a superficial region which usually gives a high background signal (final size: 290 × 290 × 20 μm^3^). All recipient cells were detected following binarization using adaptive thresholding (sensitivity = 0.01) and filtering (300 px^2^ < Area < 6000 px^2^ and eccentricity < 0.9) from the summed image of AF568 and AF647 channels for each axial slice, resulting in a 3D label matrix. Detections from each slice were linked and indexed to yield a cell-based label matrix. Background signals for each channel were measured outside of the cell label matrix. Accordingly, transfer signals in each cell were defined as the maximum of SNRs from the cell measured for each slice. The center of mass for each cell was recorded as its location.

To measure the distance of recipient cells from GFP dendrites, dendritic projections were traced from the patched cell. The GFP channel slice images were binarized, linked, and indexed as described above but using gray thresholding instead (Otsu’s threshold × 2). From the resultant label matrix, the indexes for the patched cell and not-automatically connected parts of dendrites were manually selected and linked to finalize the neuronal mask matrix. The distance of recipients from the patched soma was measured as the distance between centers of masses. The distance from the dendrites was measured by increasing the mask for the patched neuron by one pixel at each iteration and checking any contact with the recipient cell labels.

### Computational modeling of Aβ transport through dendritic DNT

Nanotubular transport of Aβ peptides between two cells connected by a DNT was modeled to explain the result of Aβ propagation from direct infusion by whole-cell patch clamping using a MATLAB script. Two cells were idealized as identical spheres (*r* = 5 μm). One of the cells was designated as a donor cell, which corresponds to the Aβ-infused cell from the experiment, with the fixed concentration of Aβ (*C*_*D*,0_ = 1.5 μM) for its connection to a reservoir while the other had no peptide in the beginning (**Fig. 4M**). Two cells were connected by dendrites protruding from each cell and a DNT connecting two dendrites. The DNT with a fixed length (*l* = 4 μm) was assumed to mediate directional transport between the dendrites of the donor to the acceptor at certain locations on the two dendrites (*x*_1_ and *x*_2_). Assuming passive transport along dendritic DNT as one-dimensional diffusion of Aβ peptides along dendrites and the DNT, we can derive the concentration of the peptides at a specific location on a dendrite *C*_*dend*_(*x*,*t*), from Fick’s second law of diffusion under the given boundary condition:

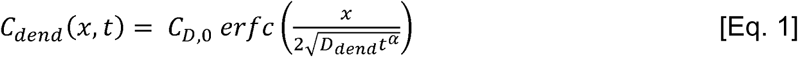

where *erfc* notes the complementary error function, *D_dend_* the one-dimensional diffusion coefficient of Aβ peptide in dendrites (∼ 55.7 μm^2^/s, calculated considering Stokes radius for the Aβ monomer from the relation of molecular weights and hydrodynamic radius of PEG) (*69, 70*), and α the anomalous exponent. Anomalous diffusion of the peptides in spiny dendrites of pyramidal neurons was hypothesized (α = 3 when the density of dendritic spines ∼ 1 ea/μm) (*69*). Considering the diffusion continues through the DNT and the connected dendrite to reach the acceptor cell, we approximate *C_NB_*(*t_i_*), the concentration at the end of the DNT, and *C_A_*(*t_i_*), the Aβ concentration in the acceptor, by the next iterative relation.

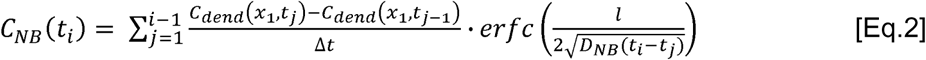

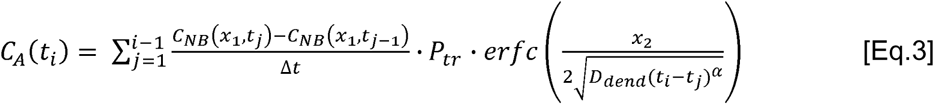

*D_NB_* describes the one-dimensional diffusion coefficient of Aβ peptide in DNTs. Passive transfer rate *P_tr_* is adopted to describe the ratio of concentrations across membranes; for complete separation without channels, *P_tr_* = 0; for complete connection by membrane fusion, *P_tr_* = 1; for conditional transfer by cellular mechanism, 0 < *P_tr_* < 1.

The quantity of active amyloid transport is simply represented by the effective flux *v*, the number of peptides transferred per unit time interval (Δ*t* = 1 sec). During the duration of transfer *t*, the amount of transported Aβ is

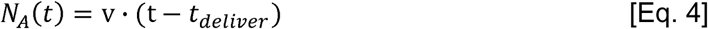

where

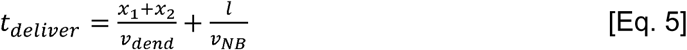

for *v_dend_* and *v_NB_* note the speed of transport on dendrites and DNTs respectively (0.75 and 0.29 μm/s assuming Kinesin-1 and Myosin-V involvement)(*71, 72*).

This preliminary model was tested to estimate values for modeling parameters. For each trial, we randomly allocated the position of DNT on dendrites (10 μm < *x*_1_ and *x*_2_ < 150) and the for a random duration (0 sec < τ < 1800 sec). From 2,000 trials, the normalized probability duration of transfer. During 30 minutes of observation, we assumed that DNTs mediate transfer distribution of *C_A_*(*t* = 1800 *sec*) was obtained supposing (1) passive (diffusive) transport only or (2) both passive and active transports. The results with differently set values of parameters were compared to fit our experimental data and yield the best parametric setting.

### Live-cell imaging of Aβ transport in dissociated neuronal culture

Cortical neurons cultured in 24-well glass bottom plates were used for the experiment when reached DIV 11-12. Our home-built fluorescent microscope equipped with a Stage Top Incubator is used for imaging while maintaining temperature (37°C) and CO_2_ concentration (5%). Cells were exposed to 1 μM of HiLyte™ Fluor 647-labeled Aβ1-42 (MW 5.5 kDa, Anaspec, Fremont, CA, USA, Cat.#: AS64161) dissolved in culture medium (> 30 min), washed twice, and imaged in phenol red-free Neurobasal Medium (1% FBS). Dynamics of Aβ peptides in neurons were imaged every second (acquisition time = 100 ms) for 5 to 10 minutes by 60X objectives (NA=1.3). Obtained image stacks were processed by Richardson-Lucy deconvolution algorithm as previously described above. ImageJ is used to select neurites of interest, draw kymographs of Aβ peptides, and measure averaged speed and processivity of each transport.

### Aβ exposure on dissociated cortical neuron

Half of the culture media was exchanged the day before sample collection with Neurobasal Media (1% FBS, 2% B27) containing 1 μM, 4 μM, or no human Aβ1-42 or scrambled Aβ1-42. After 24 hours, cells were fixed as previously described for SRRF imaging. For validation of Aβ accumulation in cells, Aβ peptides were stained by anti-Aβ (1:200; clone 4G8, BioLegend, Cat.#: 800710) and Goat anti-Mouse IgG-AF488 (Thermo Fisher Scientific, Cat.#: A11029) following the previously described procedure. For the primary antibody also detects Aβ within APP prior to cleavages, immunostained signals are labeled as Aβ/APP. Aβ accumulation in cells and other fluorescent signals were imaged using the 10× low magnification objective and selectively reflected LED lights, stitched (1535 by 1535 μm^2^, Metamorph), and analyzed using ImageJ. The boundaries of cells were first marked using binarization (Otsu’s thresholding) of the DAPI channel image and filtered (Analysis Particle, size: 20 - 200 μm^2^, circularity: 0.2 – 1). Then, we measured Aβ signals in each cell’s ROI from the Aβ image normalized by the median. For counting of DNTs, DNTs were stained and analyzed from SRRF images as described above in Sample preparation for SRRF imaging of cultured cortical neuron and Characterization of DNT from SRRF images of dissociated neurons.

#### Validation of Intraneuronal Aβ staining by the clone of antibody

Dissociated neurons were exposed to 2 μM of HiLyte™ Fluor 555-labeled Aβ1-42 (MW 5.5 kDa, Anaspec, Cat.#: 60480) to induce internalization, fixed with 4% PFA, and immunostained for 6 hrs with either Anti-Aβ Clone 4G8 (Biolegend, Cat.#: 800109) or 6E10 (Biolegend, Cat.#: 803004) followed by 1 hr incubation with Goat Anti-Mouse-IgG AF647 (Thermo Fisher Scientific, Cat.#: A21241). Images were analyzed by segmenting cells from the DAPI channel (adaptive thresholding), measuring each immunostained signal in signal-to-noise ratio from cells and the rest as background, and comparing the SNRs to decide the brightest signal using Matlab.

### Analysis of intracellular Aβ and DNT formation in AD model mouse brain

ImageJ was used to quantify intracellular Aβ signals in the somas of GFP-expressing neurons from dSRRF images, which were acquired and reconstructed as described above. For the primary antibody (clone 4G8) also detects Aβ within APP prior to cleavages, immunostained signals are labeled as Aβ/APP. A subset of the image stack with soma of interest was selected depending on the axial range of the soma and projected to obtain an averaged image. From the averaged image, the intracellular amyloid signal was measured inside the boundary of soma marked in the GFP channel manually and the background signal was measured in the remaining region for the calculation of signal-to-noise ratio. To investigate DNT formation related to intracellular Aβ accumulation, 2∼4 dendrites from each soma were tracked and analyzed as described above (morphological analysis of *in situ* DNT).

### Simulation of active Aβ propagation through dendrite-DNT network in tissue

Based on the computational model of Aβ transfer through DNTs, we simulated Aβ propagation by a DNT network in tissue. A lattice of Identical cells (N = 1,668) with a given average distance (*d* = 14.36 μm, decided from NeuN-stained images of layer 5 mPFC slices, **Supplementary Fig. 6A**) was introduced (**Supplementary Fig. 6B**, 11 × 11 × 15 cells in the volume of 158 × 158 × 115 μm^3^). We hypothesized that the probability that two cells *i* and *j* in the lattice were connected by DNTs from cell *i* is proportional to the volume of the intersection of two spheres which are centered at the position of each cell with radius *R* = 150 μm, the length of dendrites, by geometry (**Supplementary Fig. 6C**) (*62*).

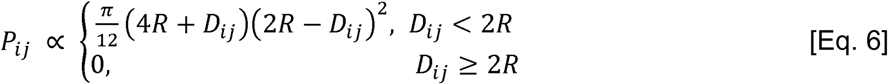

*D_ij_* corresponds to the distance between cells *i* and *j* in the lattice. The number of active DNTs for cells in the lattice is set to follow a normal distribution with simulation parameters mean *μ_NB_* and standard deviation *σ_NB_*. The number was adjusted for cells located on the boundaries by reducing it proportionally to the number of reachable cells. Now we can define a connectome matrix *M_conn_* after assigning cell pairs connected by a random permutation weighted by Eq. 6:

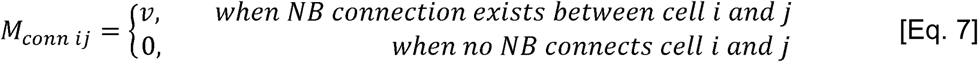

where its row-wise summation gives the total export from each cell in a unit time interval and its column-wise summation denotes the total import to each cell.

The initial number of Aβ peptides in each cell, *N_i_*(*t*), was given by an input depending on the purpose of experiments and the total Aβ number was fixed during simulation. For every time interval (Δ*t* = 1 min), *N_i_*(*t*) was updated following *M_conn_*. When a cell does not possess enough peptides to export, DNTs formed from the cell are randomly selected in order and neutralized by modifying *M_conn_* to avoid a negative amount of the peptides which is improbable, and the expected change in the system was recalculated. This adjustment is repetitively performed until none of the cells show negative for the given time point.

### Statistics

Statistical analyses were performed using MATLAB R2022a (Mathworks). All statistical tests are described in the corresponding figure legends. T-test and ANOVA were used when the normality of data was identified by Kolmogorov-Smirnov test. A *P* value of 0.05 or less was considered statistically significant.

## Figures

All the figures and schematics were created with Adobe Illustrator CS6 and BioRender.com.

## Data availability

Data that supports the findings of this study are available from the corresponding author upon reasonable request.

## Code availability

Custom MATLAB scripts used in this study are available from the corresponding author upon reasonable request.

---

**Supplementary Figure 1.**
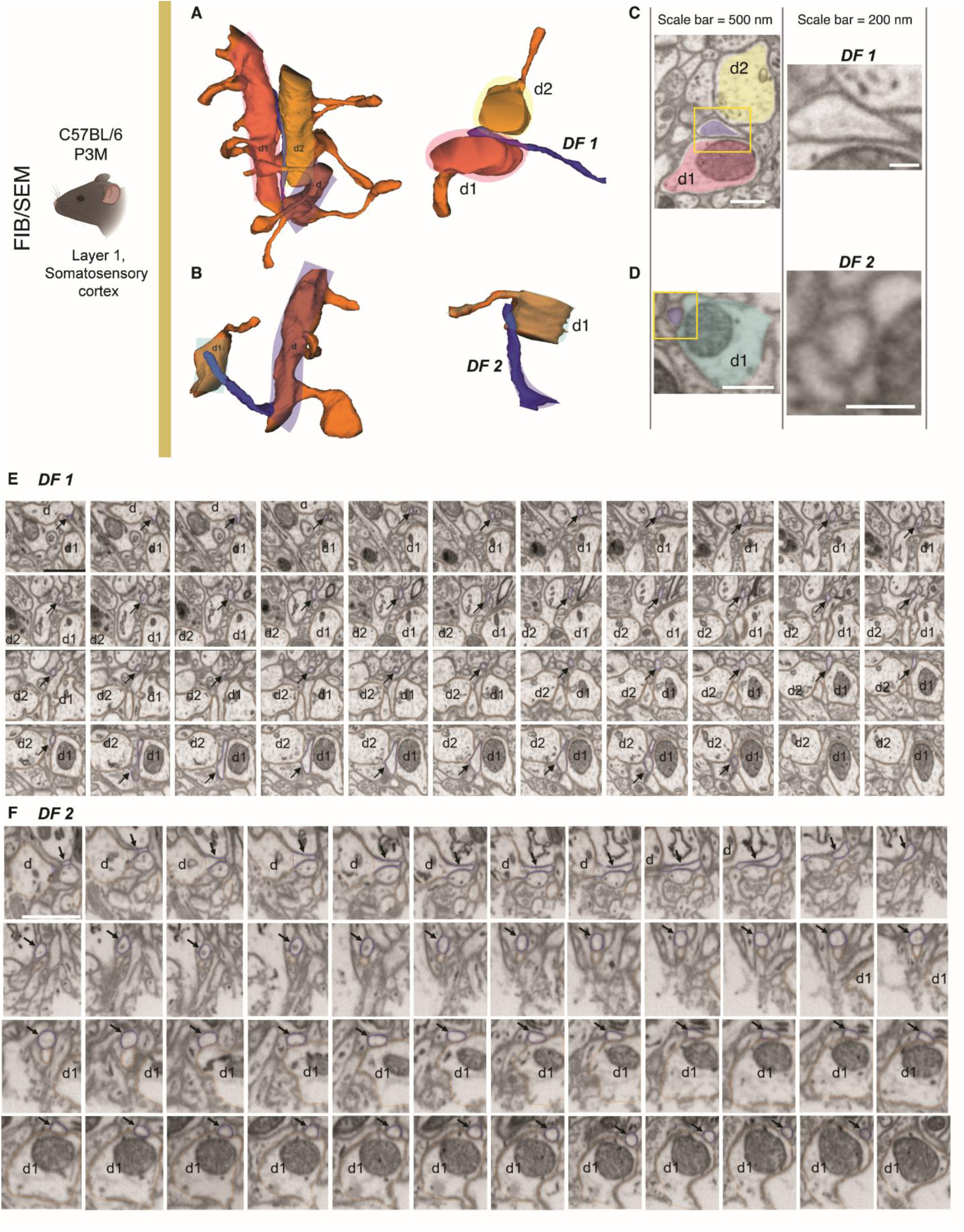
Nonsynaptic dendritic filopodia contacting other dendrites in the EM-resolved mouse brain. **(A-B)** Representative images of dendritic filopodia (DF1 and DF2) reconstructed from FIB/SEM imaging which reveal their parent dendrite (marked as d) and contacting dendrites (marked as d1 and d2; layer 1, somatosensory cortex, P3M C57BL/6). **(C-D)** Single-sectional images of filopodial terminals ((C) for DF1, (D) for DF2) contacting to the other dendrites show membrane-membrane bindings but no synaptic density. **(E-F)** Serial axial plane analysis of two dendritic filopodia represents no synaptic density along the whole contacting ranges ((E) for DF1, (F) for DF2). Arrows denote the position of filopodia. Axial step distance = 40 nm. Scale bars = 1 μm.

**Supplementary Figure 2.**
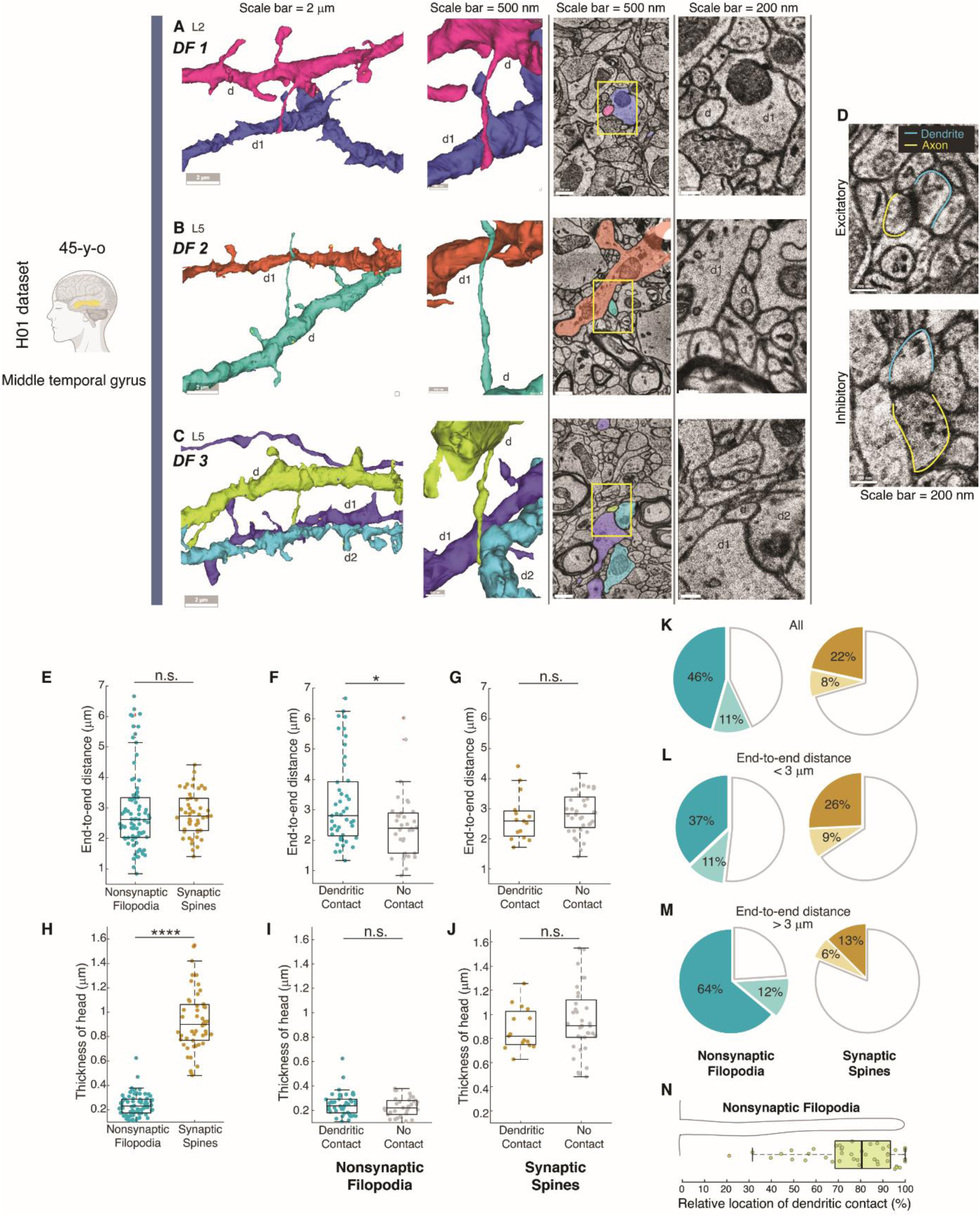
Nonsynaptic dendritic filopodia contacting other dendrites in the EM-resolved human brain. **(A-C)** Examples of dendrite-contacting filopodia without annotated excitatory or inhibitory synapses or synaptic density from layers 2 (DF1) and 5 (DF2 and DF3) of the shared H01 dataset (1^st^ column = wide view of reconstructed dendrites; 2^nd^ column = close view of contacting filopodium; 3^rd^ column = sectional image of filopodial contact; 4^th^ column = magnified sectional image of filopodial contact terminal from the yellow box). Parent dendrites = marked as d. Contacting dendrites = marked as d1 and d2. Colored points indicate annotated synapses (yellow = excitatory, blue = inhibitory). See **Supplementary Fig. 3** and **Supplementary Movies 1-3** for serial axial plane analysis. **(D)** Representative sectional images of excitatory and inhibitory synapses from the same data set showing clear synaptic densities. **(E)** End-to-end distances of non-synaptic dendritic filopodia (n = 79) and synaptic spines (n = 51) from the dendrites of layer 5 neurons. *P* = 0.56 by two-tailed t-test with equal variances. **(F)** End-to-end distances of non-synaptic dendritic filopodia with contacts to the other dendrites (n = 45) and without the contact (n = 34) from the dendrites of layer 5 neurons. *P* = 0.014 by two-tailed t-test with equal variances. **(G)** End-to-end distances of synaptic spines with contacts to the other dendrites (n = 15) and without the contact (n = 36) from the dendrites of layer 5 neurons. *P* = 0.48 by two-tailed t-test with equal variances. **(H)** Head thicknesses of non-synaptic dendritic filopodia (n = 79) and synaptic spines (n = 51) from the dendrites of layer 5 neurons. *P* = 3.5 × 10^-44^ by two-tailed t-test with equal variances. **(I)** Head thicknesses of non-synaptic dendritic filopodia with contacts to the other dendrites (n = 45) and without the contact (n = 34) from the dendrites of layer 5 neurons. *P* = 0.27 by two-tailed t-test with equal variances. **(J)** Head thicknesses of synaptic spines with contacts to the other dendrites (n = 15) and without the contact (n = 36) from the dendrites of layer 5 neurons. *P* = 0.33 by two-tailed t-test with equal variances. **(K-M)** The percentages of all investigated (K), short (L, end-to-end distance < 3 um), and long (M, end-to-end distance > 3 um) dendrite-contacting filopodia (left) and spines (right). Dark teal and brown = contact to dendritic shaft. Light teal and brown = contact to dendritic spine. **(N)** Relative locations of dendritic contacts along the lengths of nonsynaptic filopodia (n = 45; median = 81%, mean = 77% when the bottom = 0% and terminal = 100%). In the box plots, the midline, box size, and whisker indicate median, 25-75^th^ percentile, and ±2.7σ, respectively.

**Supplementary Figure 3.**
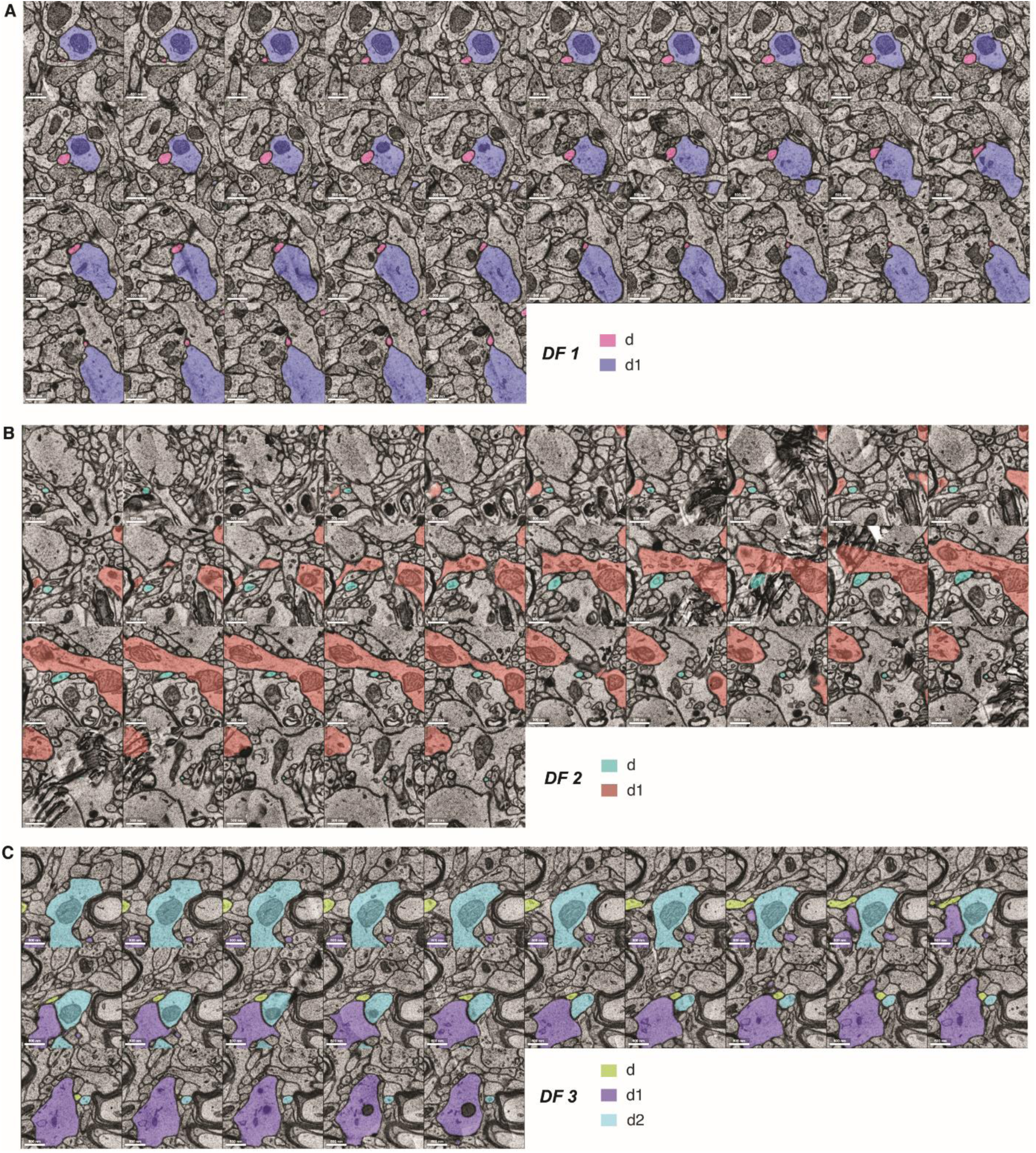
Serial axial plane analysis of nonsynaptic dendritic filopodia contacting other dendrites in the EM-resolved human brain. Serial axial plane analysis of representative dendritic filopodia represents no synaptic density along the whole contacting ranges ((A) for DF1, (B) for DF2, and (C) for DF3 of **Supplementary Fig. 2** and **Supplementary Movies 1-3**). Axial step distance = 33 nm. Parent dendrites = marked as d. Contacting dendrites = marked as d1 and d2. Scale bars = 500 nm.

**Supplementary Figure 4.**
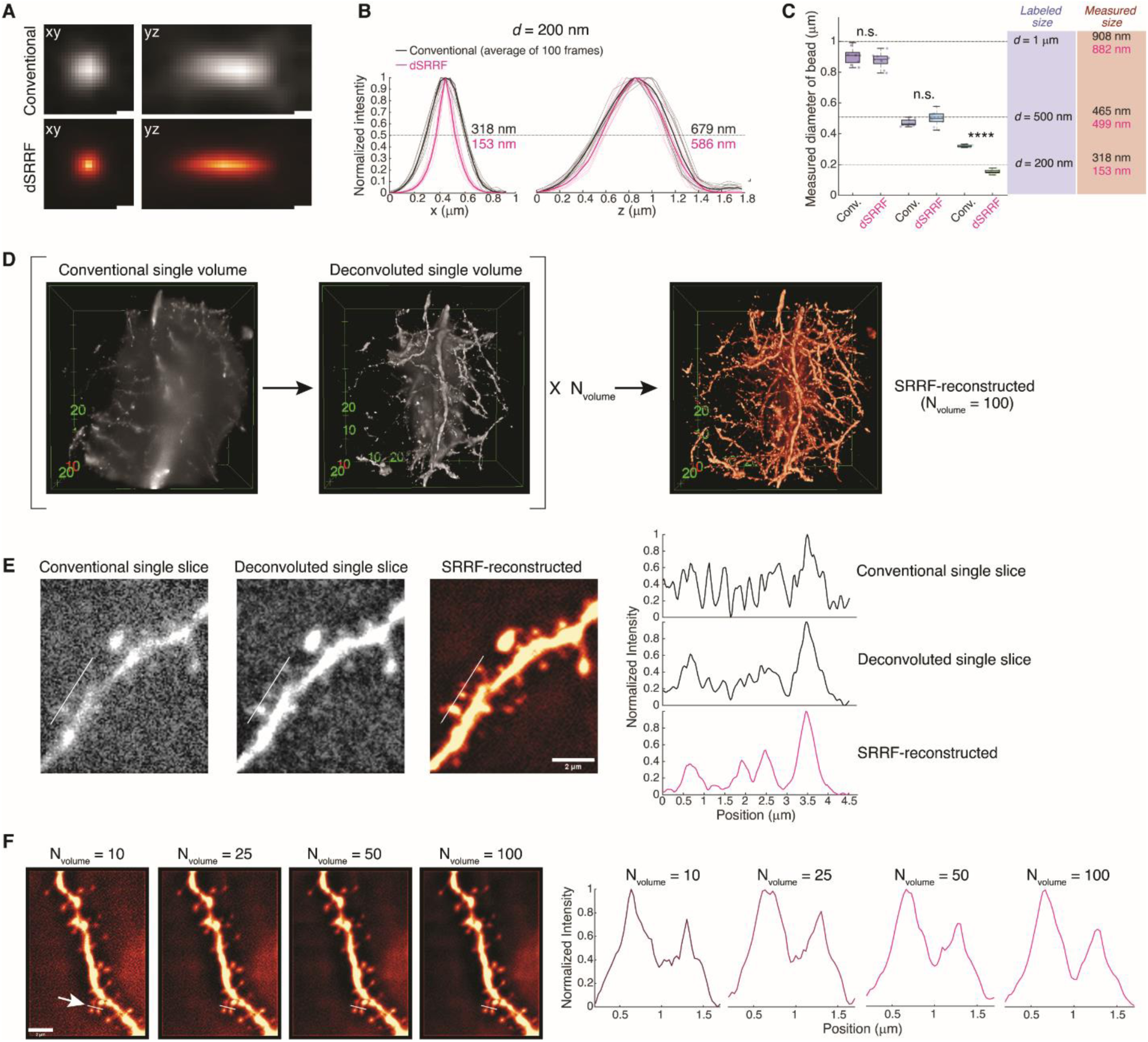
Super-resolution imaging of dendritic structure by dSRRF. **(A)** Example images of point spread function (PSF) from a fluorescent bead (diameter = 200 nm) in conventional inclined illumination microscopy (top) and dSRRF microscopy (bottom). N_volume_ for SRRF process = 100. **(B)** Comparison of the PSFs in conventional (black) and dSRRF (magenta) microscopy imaging (n = 10). The full-width-half-maximum (FWHM) of the intensity profiles represents the median diameter of the beads in measurement. Thin line = individual profiles from each bead. Thick line = the median of all profiles. **(C)** Measured diameter of beads with different sizes in volumetric scanning (axial step size = 200 nm) with conventional (black) and dSRRF (magenta) microscopy (d = 1 μm, 500 nm, and 200 nm; n = 9, 10, and 10 for each type of beads). *P* = 0.25 for d = 1 μm; *P* = 0.09 for d = 500 nm; *P* = 2.4 × 10^-18^ for d = 200 nm by two-tailed equal variances t-tests. **(D)** Representation of dSRRF processing for an image volume (65 × 65 × 40 μm^3^) obtained from the tissue-cleared brain slice of Thy1-GFP-M mouse (Supplementary Fig. 3). N_volume_ image stacks after deconvolution were used to reconstruct the volume in super-resolution (Unit for ticks in the images = μm). **(E)** Example of enhanced dendritic protrusion imaging in dSRRF. **(F)** The quality of dSRRF with increased N_volume_. n.s., not significant; **** *P* < 0.0001; In the box plots, the midline, box size, and whisker indicate median, 25-75^th^ percentile, and ±2.7σ, respectively. Scale bars = 200 nm for (A), 2 μm for (E, F)

**Supplementary Figure 5.**
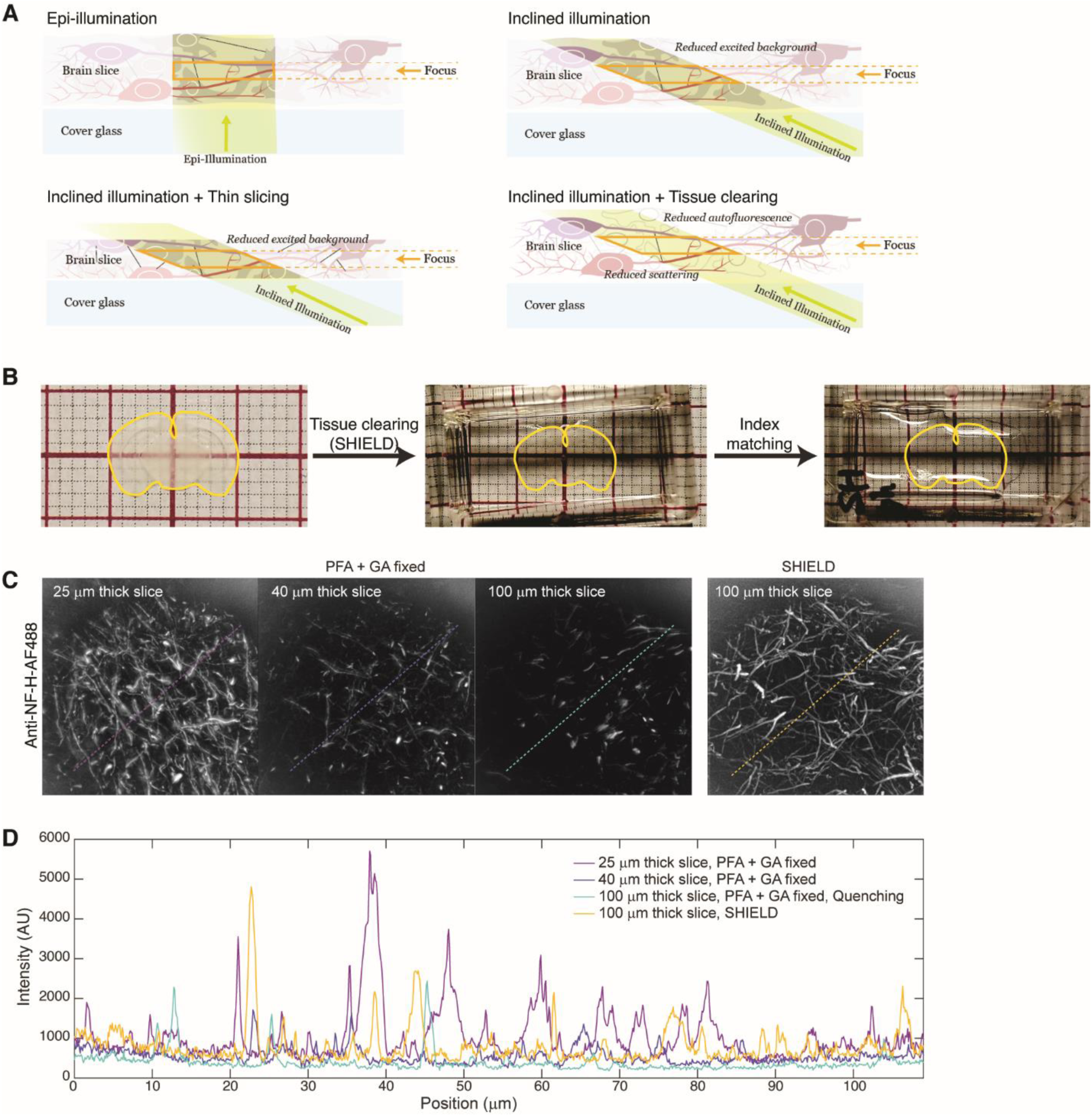
Sample preparation for dSRRF imaging of mouse brain slice. **(A)** Illustrations to explain fluorescent signal enhancement by inclined illumination, thin brain slicing, and tissue clearing. **(B)** Tissue clearing of mouse brain slice following SHIELD protocol. **(C-D)** Enhanced immunostaining signal from wild-type mouse brain slices (Anti-NF-H-AF488) by thin-slicing and tissue clearing (SHIELD).

**Supplementary Figure 6.**
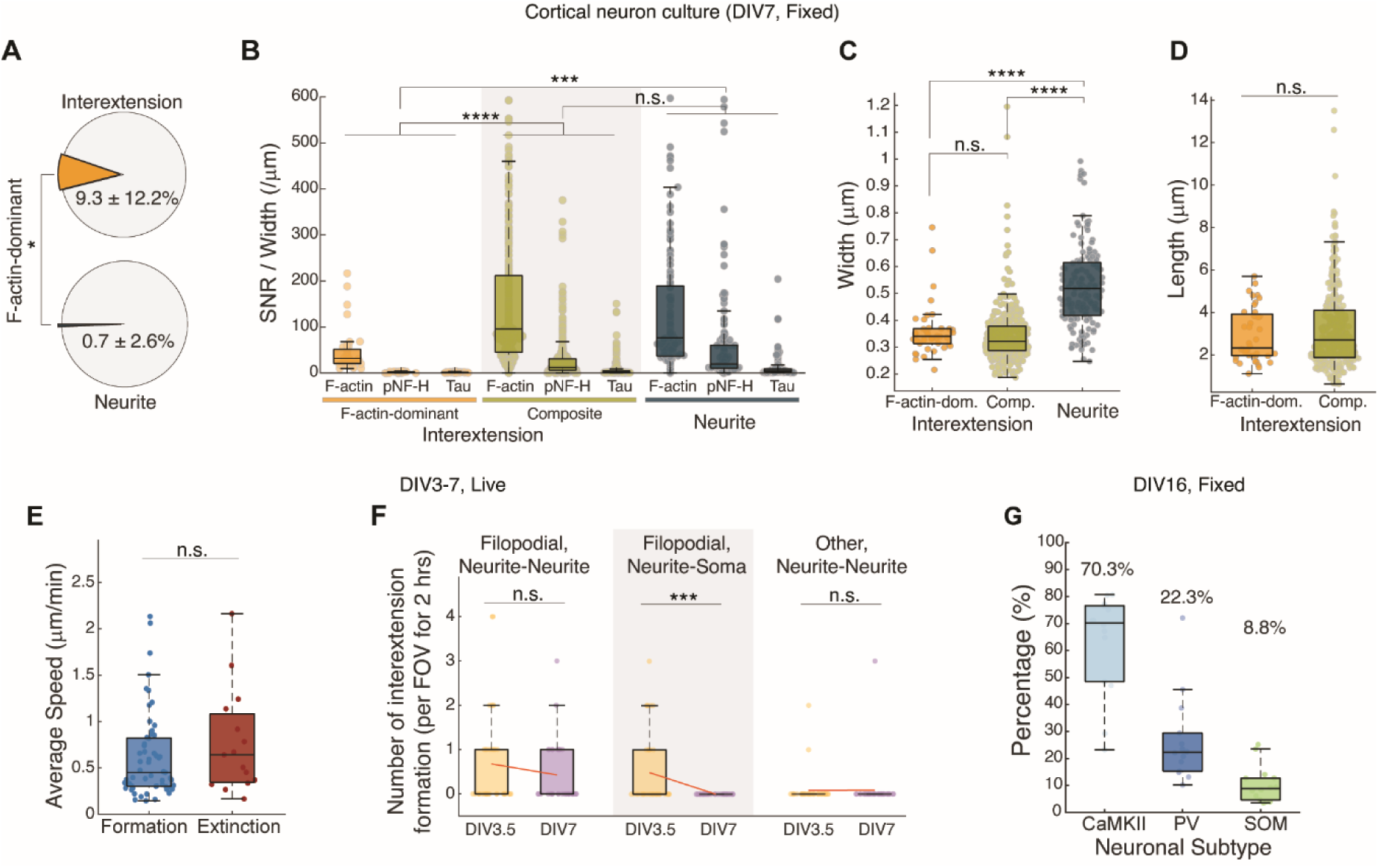
DNT formation in dissociated cortical neurons. **(A)** Percentage of F-actin-only structures in interextensions and neurites (n_FOV_ = 15) whose SNR for pNF-H and tau < 1.2 from DIV7 cortical neurons in dissociated culture (two-tailed unequal variances t-test, *P* = 0.018). **(B)** SNR of averaged intensity profile along structures, normalized by width for each cytoskeletal component from F-actin-only interextensions (left, n = 40), composite interextensions (middle, n = 351), and neurites (right, n = 144). *P* = 9.4 × 10^-5^ for Actin-IB vs. Comp-IB, *P* = 2.7 × 10^-4^ for Actin-IB vs. Primary, and *P* = 0.99 for Comp-IB vs. Primary by unbalanced two-way ANOVA, *P* = 1.4 × 10^-4^ among groups. **(C-D)** Width and length of structures (*P* = 0.77 for Actin-IB vs. Comp-IB, *P* = 6.0 × 10^-14^ for Actin-IB vs. Primary, and *P* = 0 for Comp-IB vs. Primary by one-way ANOVA; *P* = 8.8 × 10^-42^ for (C), *P* = 0.21 for Actin-IB vs. Comp-IB by two-tailed unequal variances t-test for (D)). **(E)** Average speeds of formation or extinction were obtained by dividing the length of filopodial bridges with the formation/extinction time (n = 57 and 15; *P* = 0.34 by two-tailed unequal variances t-test). Extinction events that are not clearly distinguished from photobleaching were excluded in analysis. **(F)** The number of interextension formations per FOV (94 × 94 μm^2^) for 2-hour-long time-lapse imaging. *P* = 0.25 for Filopodial N-N; *P* = 9.9 × 10^-4^ for Filopodial N-S; *P* = 0.96 for Other N-N (formation during separation of contacting neurites) by two-tailed equal variances t-tests. n_exp_ = 37 and 35 for DIV3.5 and DIV7 respectively. **(G)** Neuronal population measured by immunohistochemistry in dissociated culture of cortical neurons (n_FOV_ = 14 from 2 independent experiments, FOV = 420 X 420 μm^2^). The SNRs of immunostaining for CaMKII, PV, and SOM were compared for each cell, and the brightest signal was regarded to represent the subtype of the cell. The numbers on the bars indicate the medians. n.s., not significant; * *P* < 0.05; ** *P* < 0.01; *** *P* < 0.001; **** *P* < 0.0001; In the box plots, the midline, box size, and whisker indicate median, 25-75^th^ percentile, and ±2.7σ, respectively. Bonferroni correction followed all the ANOVA tests.

**Supplementary Figure 7.**
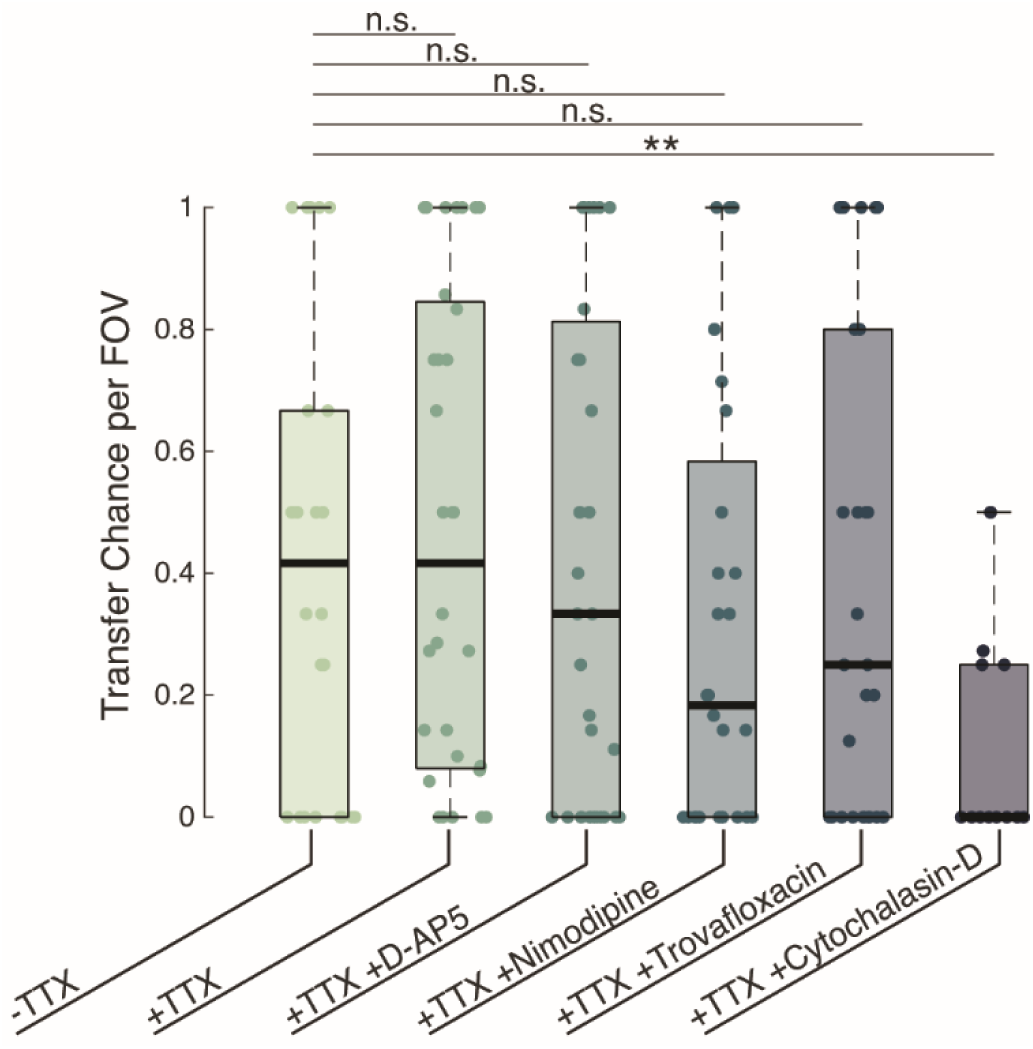
Transfer of uncaged Ca^2+^ is nonsynaptic but via DNT. To investigate the mechanism of calcium propagation to neighboring neurons, the alterations in calcium transfer chance per field of view (the portion of neighbors representing increased calcium indicator signal after uncaging) were monitored with various pharmacological inhibitions; 1 μM tetrodotoxin (TTX, sodium channel inhibitor), 100 μM D-AP5 (NMDA receptor antagonist), 20 μM Nimodipine (L-type calcium channel inhibitor), 30 μM Trovafloxacin (pannexin-1 inhibitor), and 1 μM Cytochalasin-D (nanotube formation inhibitor). *P* = 0.65, 0.86, 0.36, 0.72, and 0.002 by two-tailed unequal variances t-tests. n_FOV_ = 24 (-TTX), 32 (+TTX), 31 (+TTX +D-AP5), 28 (+TTX + Nimodipine), 29 (+TTX +Trovafloxacin), and 12 (+TTX +Cytochalasin-D) from 3 independent samples. n.s., not significant; ** *P* < 0.01; In the box plots, the midline, box size, and whisker indicate median, 25-75^th^ percentile, and ±2.7σ, respectively.

**Supplementary Figure 8.**
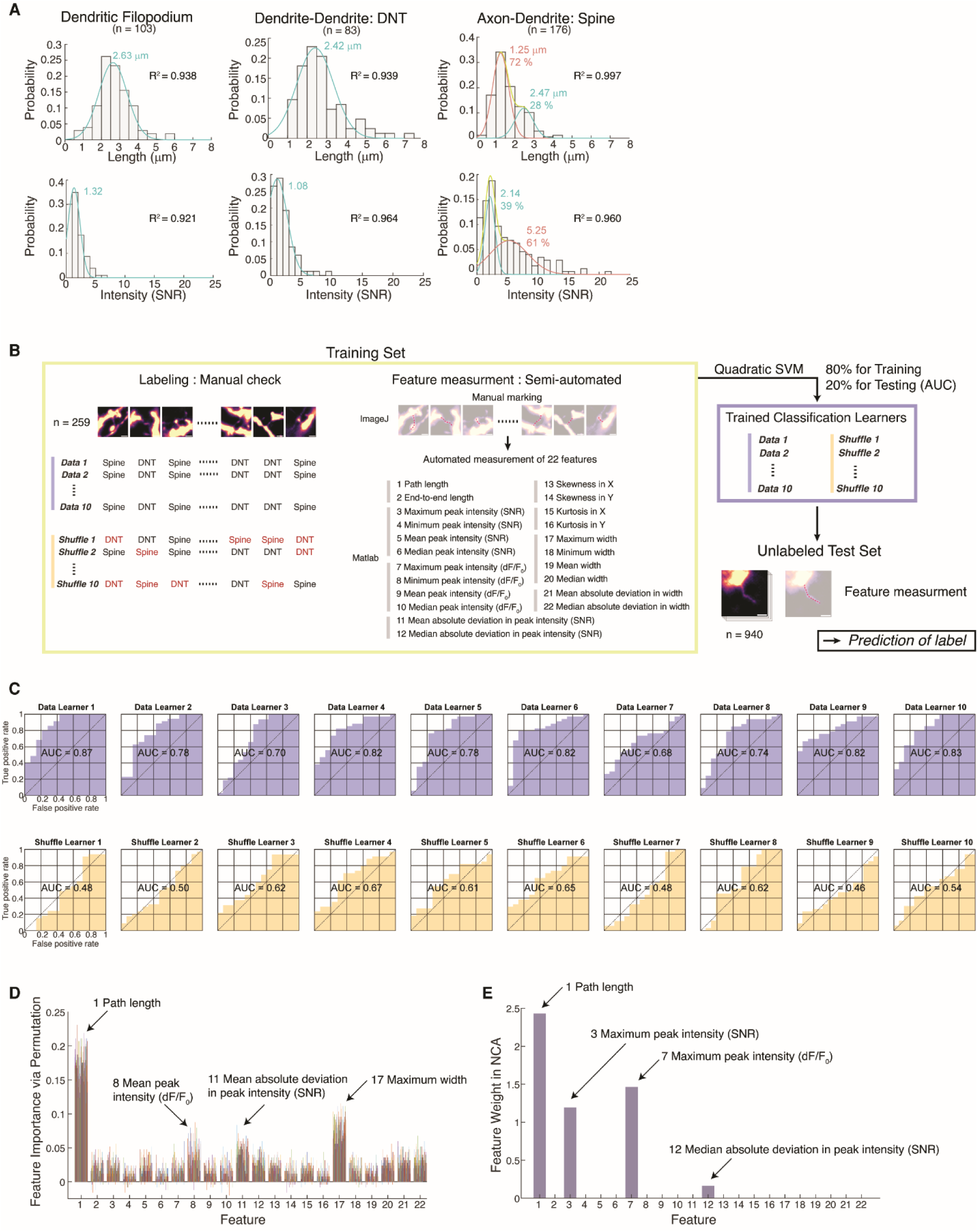
Supervised training of DNT classification learner. **(A)** Normalized probability distribution for lengths and median intensities of Filopodia, DNTs, and Spines. Curves were fitted by Gaussian with one terms (Filopodia and DNTs) or with two terms (Spines). **(B)** Training of supervised classification learner (quadratic SVM) for DNT / Spine classification. Ten models were trained with 22 morphological features of pre-labeled structures (n = 259) each for the actual data set (Data) and shuffled data set (Shuffle) and compared. **(C)** Performance of trained learners (n_learner_ = 10 for Data and Shuffle groups each) measured by AUC. **(D)** The importance of each feature in classification performance, which is defined by the reduction in the accuracy when the input values for the feature is permutated (repeated 10 times for each Data learners (n_learner_ = 10), resulting 100 bars for each feature). **(E)** The feature weight in classification performance, which is measured by the neighborhood component analysis (NCA).

**Supplementary Figure 9.**
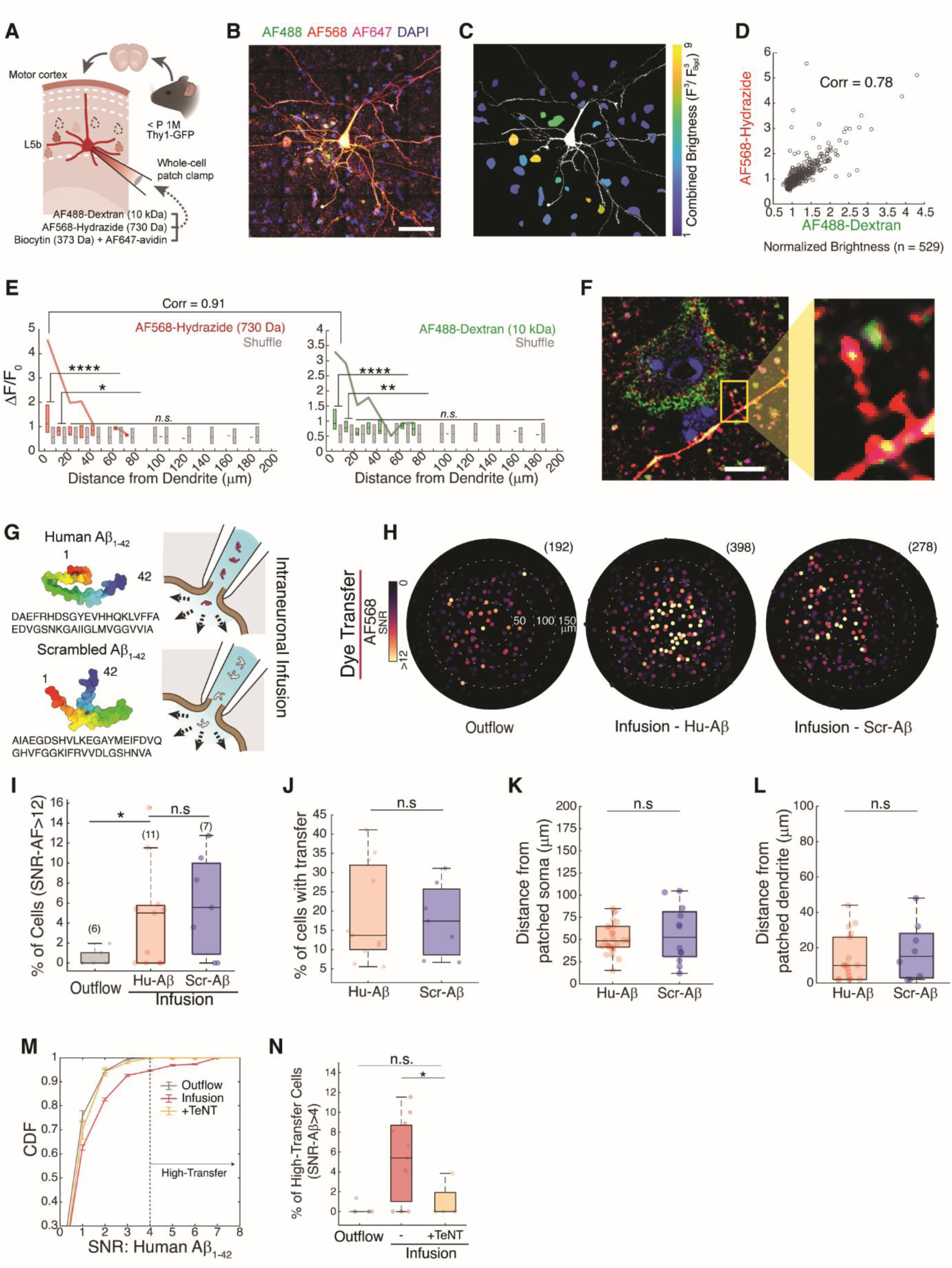
DNT-mediated molecular propagation in brain slices. **(A)** Experimental scheme of molecular propagation through DNT using patch clamping. The internal solution containing AlexaFluor568-hydrazide (730 Da; 50 μM), Dextran-AlexaFluor488 (10 kDa; 50 μM), and biocytin (0.25%w/v) was directly injected into the cell by whole-cell configuration. **(B-C)** Representative raw image of an immunostained slice (confocal microscopy, 40×, maximum projection of the 20 μm thick stack) and the outcome of automated transfer detection displayed by combined brightness (multiplication of signals from 3 fluorescence channels). **(D)** Correlation between AF568 and AF488 signals of each cell (number of cells = 529; N_exp_ = 2). The ROIs of each cell were detected automatically from the DAPI channel. Pearson’s correlation coefficient = 0.78. **(E)** Distribution of cellular fluorescent signals (ΔF/F_0_) depending on their distances from the dendrites of the patched cell and its comparison with shuffled data (shuffled match of distance and brightness). *P* < 1.0 × 10^-9^ and 0.014 for the first 2 columns of AF568 and <1.0 × 10^-7^ and 0.006 for the first 2 columns of AF488 by two-tailed unequal variances t-test. Pearson’s correlation coefficient = 0.91 when comparing lines for maximum values. **(F)** Example of transferred signal in a cell connected through a DNT from the patched cell. Scale bars = 5 μm. **(G)** Intracellular infusion of Human Aβ1-42 and Scrambled Aβ1-42 to examine the causal effect of Human Aβ1-42 in DNT-mediated molecular transport. Folding forms were predicted by AlphaFold2. **(H)** Propagation patterns of Outflow (left; n = 192), AF568 in Human (middle; n = 398), and Scrambled (right; n = 278) Aβ1-42 infusion experiments. Origin = position of the paired patched soma. **(I)** Percentage of high-transfer cells (SNR-AF > 12) from all detected cells in the imaged volume from extracellular outflow (n_exp_ = 6), Human (n_exp_ = 11) and Scrambled (n_exp_ = 7) Aβ1-42 infusion. *P* = 0.029 and 0.61 for outflow vs. Hu-Aβ and Hu-Aβ vs. Scr-Aβ respectively, by two-tailed unequal variances t-tests. **(J)** Percentage of detected cells in the imaged volume (290 × 290 × 20 μm^3^) estimated from the measured density of neurons in NeuN-stained mPFC (Supplementary Fig. 12A, ∼ 1.5 × 10^-4^ /μm^3^). *P* = 0.79 by two-tailed unequal variances t-test. **(K-L)** Locations of high-transfer cells in two groups were measured by distance from the soma (K) or distance from the dendrites (L) of the patched cell. *P* = 0.66 (n = 26 and 12 for high-transfer cells in Human and Scrambled Aβ1-42 infusion respectively) and 0.55 (n = 18 and 8) by two-tailed unequal variances t-tests. **(M)** CDFs of fluorescent signals from detected recipients by 100 nM Tetanus toxin in the internal solution (+TeNT). n_exp_ = 6, 11, and 4 for outflow, infusion, and infusion with TeNT respectively. **(N)** Fraction of high-transfer cells (SNR-Aβ1-42 > 4) from detected recipients. n_exp_ = 6, 11, and 4 for outflow, infusion, and infusion with TeNT respectively. *P* = 0.51 for outflow vs. TeNT and 0.017 for infusion vs. TeNT by two-tailed unequal variances t-tests. n.s., not significant; * *P* < 0.05; ** *P* < 0.01; *** *P* < 0.001; **** *P* < 0.0001; In the box plots, the midline, box size, and whisker indicate median, 25-75^th^ percentile, and ±2.7σ, respectively.

**Supplementary Figure 10.**
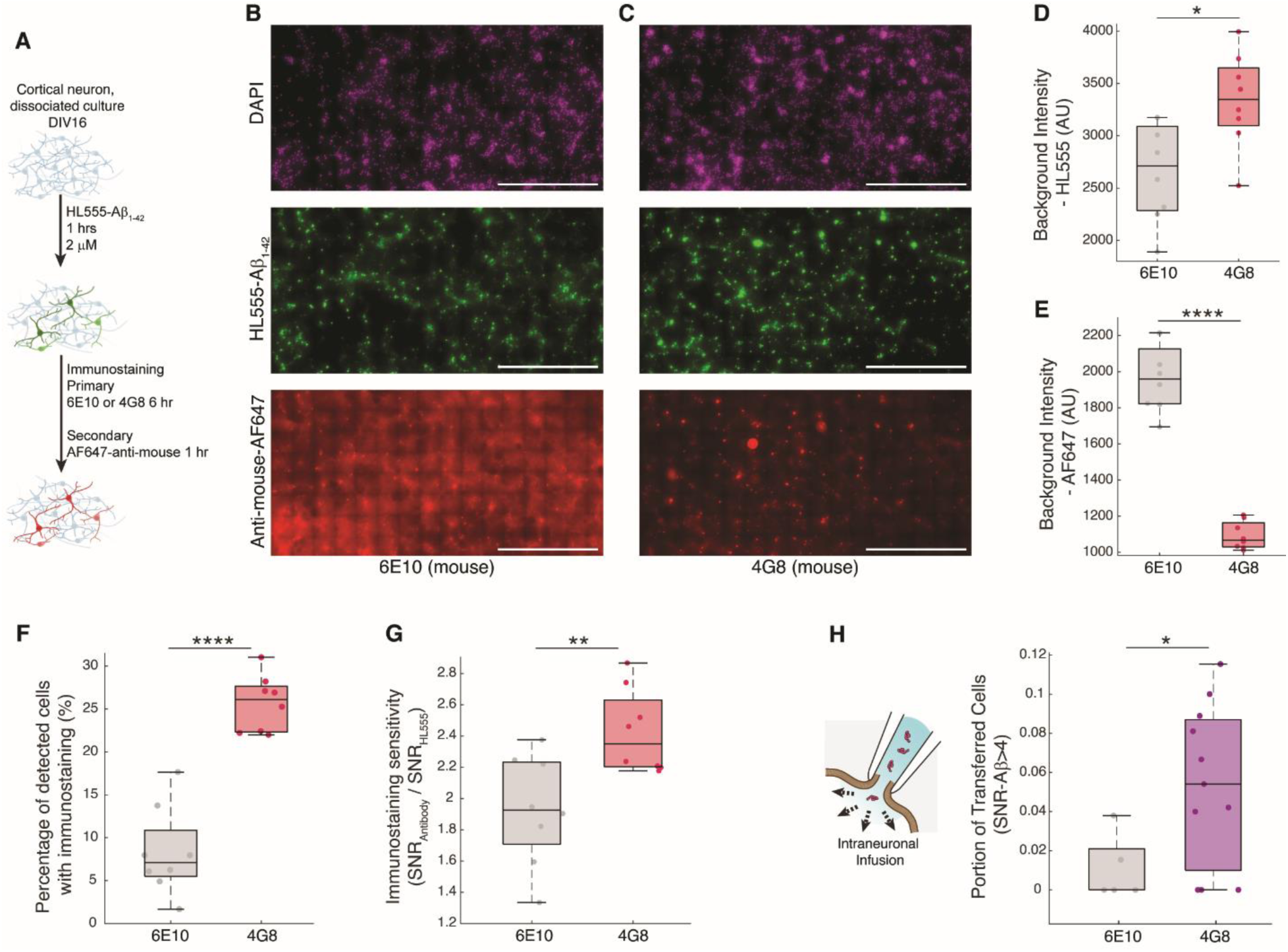
Immunostaining of intraneuronal beta-amyloid depending on the clone of antibody. **(A)** Experimental procedure for detection of Aβ1-42 in dissociated neurons. Short exposure (1 hr) to 2 μM HiLyte Fluor 555 conjugated Aβ1-42 was enough to increase intraneuronal amyloid level. Following permeablization and PFA fixation, Aβs were stained either by the clone 6E10 or 4G8 and detected with same secondary antibody (Alexa Fluor 647-anti-mouse). **(B-C)** Representative images of stained neurons in DAPI, HiLyte Fluor 555, and Alexa Fluor 647 channels respectively by clone 6E10 (B) or 4G8 (C). Scale bar = 1 mm. **(D-E)** The averaged background intensities of each ROI (1565 X 783 μm^2^; n = 8 for each clone) in the HiLyte Fluor 555 (D) and Alexa Fluor 647 (E) channels. *P* = 7.4 × 10^-9^ (D) and 0.01 (E) by two-tailed t-tests with equal variances. **(F)** The percentage of successful detection from the Alexa Fluor 647 channel while the detection from the HiLyte Fluor 555 channel were regarded as the ground truth. Unbiased detection of cells was done using adaptive thresholding. *P* = 1.2 × 10^-6^ by two-tailed t-tests with equal variances. **(G)** The sensitivity of immunostaining defined as the ratio of cellular SNRs in the Alexa Fluor 647 channel divided by SNRs in the HiLyte Fluor 555 channel. *P* = 0.0067 by two-tailed t-tests with equal variances. **(H)** The measured portion of cells with transferred Aβ in intraneuronal infusion by patch clamping (**Fig. 4**) by the clone of amyloid antibody (6E10, n = 5 or 4G8, n = 11). *P* = 0.011 by two-tailed t-tests with equal variances. n.s., not significant; * *P* < 0.05; ** *P* < 0.01; *** *P* < 0.001; **** *P* < 0.0001; In the box plots, the midline, box size, and whisker indicate median, 25-75^th^ percentile, and ±2.7σ, respectively.

**Supplementary Figure 11.**
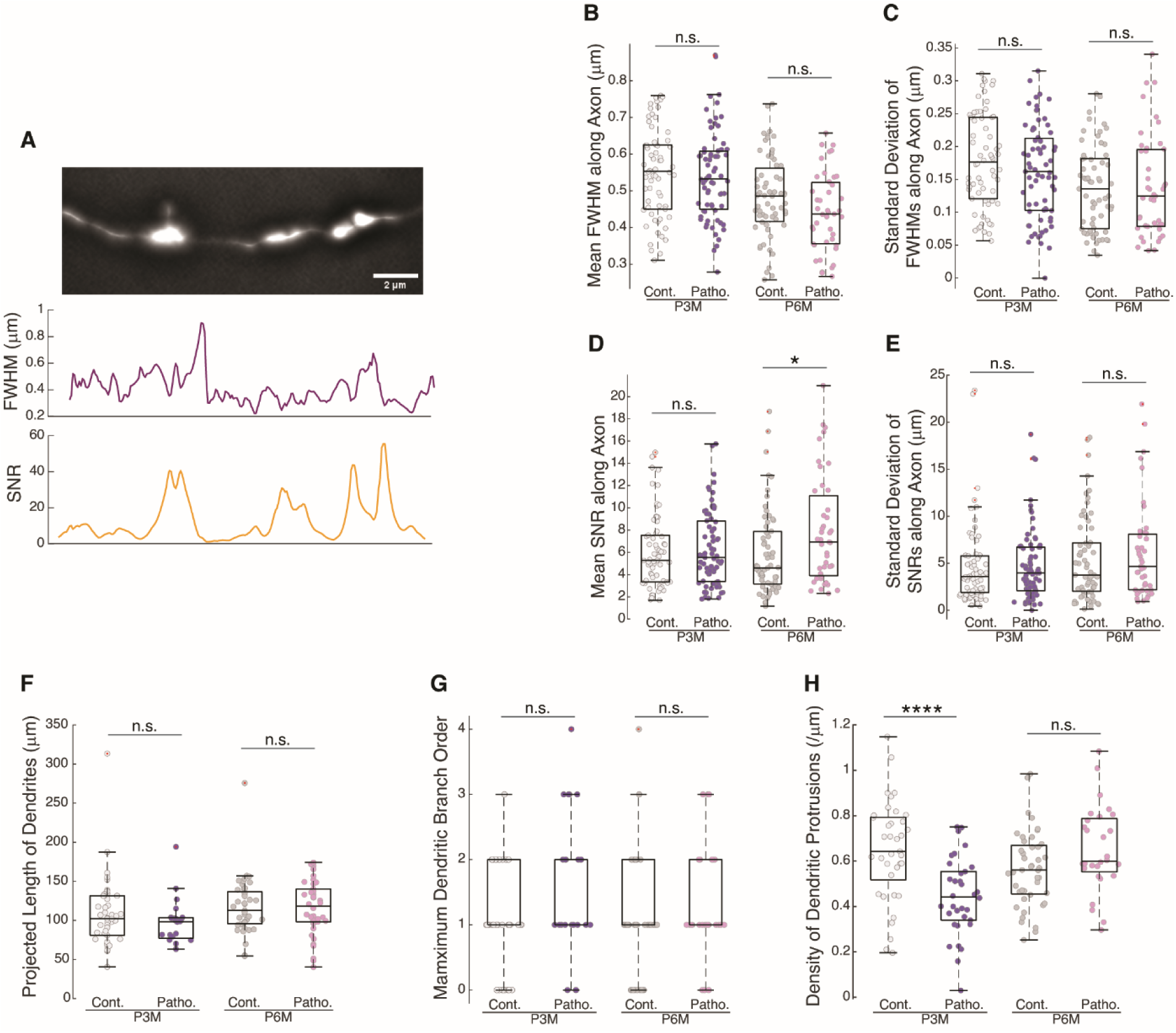
Pathological alteration of the morphologies of axons and dendrites in the amyloid pathology model brains. **(A)** An example of FWHM (full width at half maximum) and SNR (signal-to-noise ratio) detection from a segment of an axon imaged in the same field-of-view with DNTs. FWHMs can be locally inaccurate depending on the focus. **(B)** Mean FWHM along axons (n = 66, 71, 63, and 43; *P* = 1 for Cont. vs. Patho. in P3M; *P* = 0.42 for Cont. vs. Patho. in P6M; by one-way ANOVA; *P* = 1.5 × 10^-6^; 3 mice for each group). **(C)** Standard deviation of local FWHMs along axons (*P* = 0.86 for Cont. vs. Patho. in P3M; *P* = 1 for Cont. vs. Patho. in P6M; by one-way ANOVA; *P* = 0.0020). **(D)** Mean SNR along axons (*P* = 1 for Cont. vs. Patho. in P3M; *P* = 0.034 for Cont. vs. Patho. in P6M; by one-way ANOVA; *P* = 0.028). **(E)** Standard deviation of local SNRs along axons (*P* = 1 for Cont. vs. Patho. in P3M; *P* = 1 for Cont. vs. Patho. in P6M; by one-way ANOVA; *P* = 0.280). **(F)** Projected length of dendrites (from soma to the end imaged in each slice) analyzed for DNT quantification (n = 37, 18, 44, and 28; *P* = 1 for Cont. vs. Patho. in P3M; *P* = 1 for Cont. vs. Patho. in P6M; by one-way ANOVA; *P* = 0.12; 3 mice for each group). **(G)** The maximum dendritic branch order from each dendrite (n = 37, 34, 44, and 28; *P* = 0.35 for Cont. vs. Patho. in P3M; *P* = 1 for Cont. vs. Patho. in P6M; by one-way ANOVA; *P* = 0.20). **(H)** The density of dendritic protrusions along dendrites of (F) (n = 37, 34, 44, and 28; *P* = 5.8 × 10^-5^ for Cont. vs. Patho. in P3M; *P* = 0.32 for Cont. vs. Patho. in P6M; by one-way ANOVA; *P* = 1.5 × 10^-5^). n.s., not significant; * *P* < 0.05; **** *P* < 0.0001; In the box plots, the midline, box size, and whisker indicate median, 25-75^th^ percentile, and ±2.7σ, respectively. Bonferroni correction followed all the ANOVA tests.

**Supplementary Figure 12.**
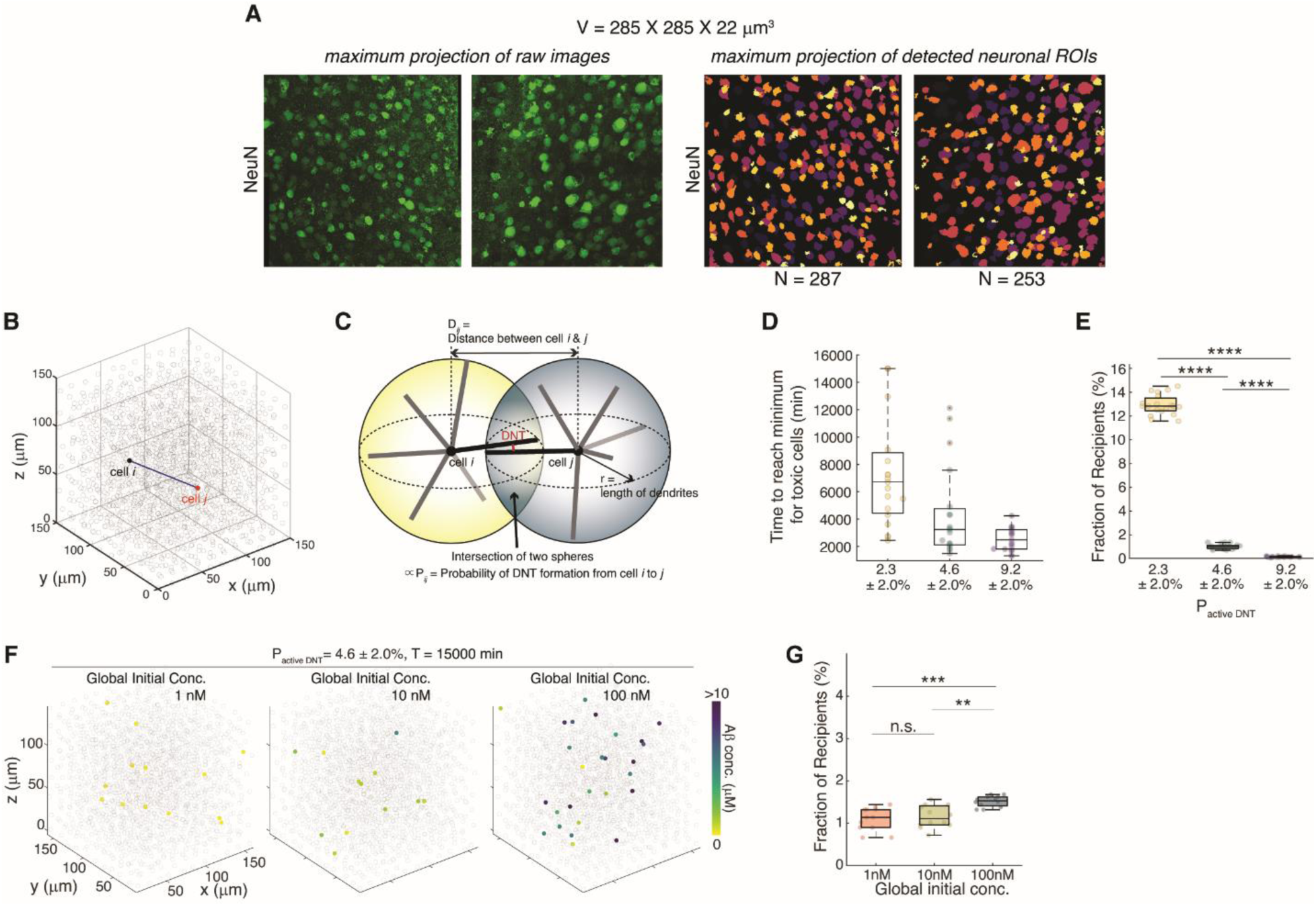
Generation of DNT network model in mPFC and simulation. **(A)** Automated detection of neurons in layer 5 mPFC from NeuN-stained brain slices to measure the density of neurons in the region (∼ 1.5 × 10^-4^ /μm^3^). **(B)** Generation of tissue model of neuronal DNT network by placing artificial neurons in a lattice (N = 1,668, 11 × 11 × 15 cells in the volume of 158 × 158 × 115 μm^3^). The average distance between cells (14.36 μm) was given from NeuN-stained images of layer 5 mPFC slices (A). **(C)** For each pair of cells in the lattice, the probability of DNT formation from cell to cell was calculated based on the intersection of two spheres that were centered on each cell with a radius of 150 μm, the maximum length of dendrites in this simulation. **(D)** Time required for toxic cells to reach the minimum Aβ concentration in burden sharing simulation of single-cell toxicity (**Fig. 6E**). n = 19 for each group. **(E)** The fraction of recipients from the whole cells in burden-sharing simulation (n = 19 for each group; *P* = 4.8 × 10^-55^ for 2.3% vs. 4.6%, *P* = 1.1 × 10^-56^ for 2.3% vs. 9.2%, and *P* = 7.3 × 10^-6^ for 4.6% vs. 9.2% by one-way ANOVA; *P* = 9.0 × 10^-59^). **(F)** Examples of aggregation in particular cells under global exposure resulting in the given initial intracellular concentrations. **(G)** The fraction of recipients from the whole cells in global exposure simulation (n = 12 for each group; *P* = 1 for 1 nM vs. 10 nM, *P* = 0.00031 for 1 nM vs. 100 nM, and *P* = 0.0017 for 10 nM vs. 100 nM by one-way ANOVA; *P* = 0.00017). In the box plots, the midline, box size, and whisker indicate median, 25-75^th^ percentile, and ±2.7σ, respectively.

**Supplementary Table.** Multi-comparison of results from ANOVA tests in DNT-mediated calcium propagation experiment.

| Groups Compared |  | *P* - Targets | *P* - Neighbors |
| --- | --- | --- | --- |
| DMSO | CBX | 1 | **1.45E-14** |
| DMSO | Cyt-D | 0.978405 | **1.94E-15** |
| DMSO | BAPTA | **2.26E-17** | **1.05E-19** |
| CBX | Cyt-D | 0.096164 | 1 |
| CBX | BAPTA | **5.44E-12** | **0.002525** |
| Cyt-D | BAPTA | **6.14E-21** | **0.000137** |

|  | *N* |  |  |  |
| --- | --- | --- | --- | --- |
| | 20-30 $\mu\text{m}$ | 30-40 $\mu\text{m}$ | 40-50 $\mu\text{m}$ | 50-60 $\mu\text{m}$ |
| DMSO | 26 | 31 | 18 | 6 |
| CBX | 36 | 58 | 22 | 7 |
| CytD | 29 | 35 | 33 | 8 |

| Time | Groups Compared |  | *P* |  |  |  |
| --- | --- | --- | --- | --- | --- | --- |
| | | | 20-30 $\mu\text{m}$ | 30-40 $\mu\text{m}$ | 40-50 $\mu\text{m}$ | 50-60 $\mu\text{m}$ |
| 10 - 20<br>sec | DMSO | CBX | 0.740904 | 0.926127 | **0.038605** | 0.713928 |
|  | DMSO | Cyt-D | 1 | 0.158547 | **0.001941** | 1 |
|  | CBX | Cyt-D | 1 | 0.708482 | 1 | 0.304039 |
| 30 - 40<br>sec | DMSO | CBX | 0.267705 | **0.006133** | 0.565421 | 1 |
|  | DMSO | Cyt-D | 0.241336 | **0.004777** | 0.493918 | **0.042258** |
|  | CBX | Cyt-D | 1 | 1 | 1 | 0.061897 |
| 50 - 60<br>sec | DMSO | CBX | **0.014232** | **7.97E-06** | **0.011678** | 0.734698 |
|  | DMSO | Cyt-D | **0.006871** | **8.57E-07** | **0.01343** | **0.006448** |
|  | CBX | Cyt-D | 1 | 0.767047 | 1 | 0.075544 |
| 70 - 80<br>sec | DMSO | CBX | **0.015618** | **9.01E-07** | **0.002364** | 0.329745 |
|  | DMSO | Cyt-D | **0.008515** | **1.01E-07** | **0.001947** | **0.004464** |
|  | CBX | Cyt-D | 1 | 0.744544 | 1 | 0.150714 |
| 90 - 100<br>sec | DMSO | CBX | **0.000402** | **7.03E-09** | **0.003035** | 0.136603 |
|  | DMSO | Cyt-D | **3.51E-05** | **1.92E-06** | **0.003214** | **0.004011** |
|  | CBX | Cyt-D | 0.712392 | 1 | 1 | 0.238996 |

## Supplementary Movies

**Supplementary Movie 1.** Serial axial planes of nonsynaptic dendritic filopodium along the whole contacting ranges to the other dendrite in the EM-resolved human brain (DF1; **Supplementary Figs. 2A and 3A**; H01 dataset). Axial step distance = 33 nm. Scale bar = 500 nm.

**Supplementary Movie 2.** Serial axial planes of nonsynaptic dendritic filopodium along the whole contacting ranges to the other dendrite in the EM-resolved human brain (DF2; **Supplementary Figs. 2B and 3B**; H01 dataset). Axial step distance = 33 nm. Scale bar = 500 nm.

**Supplementary Movie 3.** Serial axial planes of nonsynaptic dendritic filopodium along the whole contacting ranges to the other dendrite in the EM-resolved human brain (DF3; **Supplementary Figs. 2C and 3C**; H01 dataset). Axial step distance = 33 nm. Scale bar = 500 nm.

**Supplementary Movie 4.** Filopodium-originated formation of an intermediate bridge in time-lapse imaging of NeuO-stained cortical neurons in dissociated culture (DIV3.5; **Fig. 1L**). Video rate in acquisition = 1 min per frame. 4 min in experiment = 1 sec in play (Total 20 min in experiment = 5 secs in play). Scale bar = 5 μm.

**Supplementary Movie 5.** Extinction of an intermediate bridge into filopodium in time-lapse imaging of NeuO-stained cortical neurons in dissociated culture (DIV7; **Fig. 1L**). Video rate in acquisition = 1 min per frame. 15 min in experiment = 1 sec in play (Total 90 min in experiment = 6 secs in play). Scale bar = 5 μm.

**Supplementary Movie 6.** Propagation of uncaged calcium among immature cortical neurons (DIV4) following exclusive UV exposure (T = 0) on a single neuron (**Fig. 2A**). Neuronal morphology was dSRRF-resolved (gray). Calcium signal was detected using Cal-590-AM indicator (color). Acquisition rate = 100 ms per frame. 10 sec in experiment = 1 sec in play (Total 100 sec in experiment = 10 secs in play). Scale bar = 10 μm.

**Supplementary Movie 7.** Propagation of uncaged calcium through a somatic NT between immature cortical neurons (DIV4) following exclusive UV exposure (T = 0) (**Fig. 2B**). Neuronal morphology was dSRRF-resolved (gray). Calcium signal was detected using Cal-590-AM indicator (color). Acquisition rate = 100 ms per frame. 10 sec in experiment = 1 sec in play (Total 100 sec in experiment = 10 secs in play). Scale bar = 5 μm.

**Supplementary Movie 8.** Time-lapse imaging of HiLyte Fluor 647 conjugated Aβs (red) in dissociated cortical neuron (DIV11). Video rate in acquisition = 1 s per frame. 30 sec in experiment = 1 sec in play (Total 5 min in experiment = 10 secs in play). Scale bar = 10 μm.

**Supplementary Movie 9.** Active transports of HiLyte Fluor 647 conjugated Aβs (red) along a neurite of dissociated cortical neuron (DIV11) in time-lapse imaging (**Fig. 4P**). Video rate in acquisition = 1 s per frame. 30 sec in experiment = 1 sec in play (Total 5 min in experiment = 10 secs in play). Scale bar = 10 μm.

**Supplementary Movie 10.** Active transports of HiLyte Fluor 647 conjugated Aβs (red) along a DNT of dissociated cortical neuron (DIV11) in time-lapse imaging (**Fig. 4U**). The NeuO-stained DNT (cyan) was resolved from dSRRF. Video rate in acquisition = 0.5 s per frame. 30 sec in experiment = 1 sec in play (Total 2 min in experiment = 4 secs in play). Scale bar = 5 μm.

**Supplementary Movie 11.** Synchronized filopodial emergence (T = 1:14) in time-lapse imaging of NeuO-stained cortical neurons in dissociated culture (DIV3.5). Video rate in acquisition = 1 min per frame. 15 min in experiment = 1 sec in play (Total 2 hr 11 min in experiment = ∼ 9 secs in play). Scale bar = 10 μm.

**Supplementary Movie 12.** A representative simulation of toxic cell rescue by burden-sharing in a modeled DNT network (P_active DNT_ = 4.6 ± 2.0%; Fig. 6C). The color of the markers represented the Aβ concentration in each cell (gray = 0; Initial Aβ concentration in a toxic cell = 20 μM where neurotoxicity in neurons appears for 10 μM).

**Supplementary Movie 13-17.** Representative simulations of intracellular accumulation by Aβ distribution under global amyloid exposure (100 nM) in modeled DNT networks (13 for P_active DNT_ = 2.3 ± 2.0%, 14 for P_active DNT_ = 4.6 ± 2.0%, 15 - P_active DNT_ = 9.2 ± 1.0%, 16 - P_active DNT_ = 9.2 ± 2.0%, and 17 - P_active DNT_ = 9.2 ± 3.0%; Fig. 6F).

